# Neuropilin 2 stabilises adherens junctions and protects against endothelial activation by promoting the interaction between VE cadherin and p120 catenin

**DOI:** 10.1101/2025.03.09.642194

**Authors:** CJ Benwell, N Ilker, A Nicklin, LB Vaughn, M Firoglani-Moschi, CA Price, L Mitchell, T Liu, SD Robinson

## Abstract

The mechanosensing properties of endothelial cell-cell junctions are essential for vascular beds to respond to the mechanical forces exerted by blood flow. In states of disturbed flow, endothelial cells (ECs) become activated and transition to a pro-inflammatory, atheroprone phenotype. Here, we investigated the role of transmembrane glycoprotein neuropilin 2 (NRP2) in maintaining adherens junction integrity using cultured immortalised mouse ECs and a genetically modified mouse model to demonstrate the effects of an endothelial-specific deletion of Nrp2 *in vivo*. We reveal that, akin to its ortholog, Nrp1, Nrp2 exists as a constituent of adherens junctions, maintaining surface availability of VE cadherin by promoting its interaction with p120 catenin. As a consequence, endothelial knockout mice (Nrp2*^flfl^*.*EC^KO^*) display hyperpermeable retinal vasculature during development. Nrp2 depletion was subsequently found to activate key pro-inflammatory cytokines and adhesion molecules known to participate in the progression of atherogenesis, in addition to increased immune cell attachment aortic plaque development. These findings describe a role for Nrp2 in maintaining junctional signalling in ECs, protecting against endothelial activation during a state of vascular disease.

## Introduction

Endothelial cell (EC) junctions are not only critical for maintaining vascular integrity, but also for determining the semi-permeable quality of the endothelium itself, controlling the exchange of nutrients and immune cells from the circulation to the neighbouring tissue [1], [2]. During a state of cardiovascular disease, the endothelium is activated by inflammatory cytokines that disrupt the integrity of intercellular junctions, promoting vascular permeability, the trans-endothelial migration of invading leukocytes, and tissue inflammation [3], [4], [5].

Vascular endothelial (VE)-cadherin (Cdh5) is known as the principal component of EC adherens junctions, which together with armadillo adaptor proteins α catenin, β-catenin, p120-catenin and platelet endothelial cell adhesion molecule-1 (Pecam1), form a junctional mechanosensing complex that integrates and transmits mechanical force to modulate intracellular signalling in ECs [6], [7]. Adherens junctions are linked to the actin cytoskeleton via β-catenin and p120-catenin, which associate to the cytoplasmic tail of VE cadherin, enabling both junction remodelling and ensuring the plasticity of already established cell junctions. The association between p120-catenin and VE cadherin is also known to regulate VE cadherin availability at junctional adhesions by inhibiting its clathrin-mediated endocytosis (CME) from the plasma membrane and its subsequent intracellular processing. As a consequence of inhibiting this interaction, adherens junctions become destabilised and endothelial barrier function is compromised resulting from reduced endogenous VE cadherin expression [7], [8], [9], [10].

Neuropilin 2 (Nrp2) is a transmembrane glycoprotein with defined roles in ECs as a co-receptor for vascular endothelial growth factor (VEGF) and Semaphorin ligands, in addition to regulating extracellular matrix signalling by acting as an adhesion molecule [11], [12]. The more extensively studied ortholog of Nrp2, Nrp1, is also known to integrate VEGF-dependent and independent EC signalling, and to modulate vascular permeability by interacting with and regulating the turnover of VE cadherin [10], [13]. VE cadherin’s association with Nrp1, which occurs in a sheer-stress-dependent manner, has subsequently been revealed to stabilise adherens junctions by promoting the interaction between VE cadherin and p120 catenin, Nrp1 silencing inducing junction disassembly and disorganised actin remodelling. Furthermore, Nrp1 was shown to protect against endothelial activation under laminar flow, its genetic deletion from mouse endothelium increasing the expression of pro-inflammatory adhesion molecules, leukocyte rolling and atherosclerotic plaque deposition [10]. Despite these investigations however, the role of Nrp2 in regulating barrier function is poorly understood.

Akin to their shared function in regulating the intracellular transport of α5 integrin in ECs [14], we reveal Nrp2 also promotes the association between VE cadherin and p120-catenin to stabilise adherens junctions and the actin cytoskeleton. As a result, Nrp2 depletion stimulates VE cadherin endocytosis and its lysosomal degradation. Endothelial Nrp2 knockout mice exhibited an abnormally diffuse VE cadherin distribution in the postnatal retina, resulting in a high incidence of junctional gaps and vascular leakage. Transcriptomic profiling of Nrp2 depleted ECs revealed an upregulation of genes associated with vascular inflammation. *In vitro* and *in vivo* Nrp2 silencing was found to result in elevated inflammatory-associated adhesion marker expression and monocyte-endothelium attachment, in addition to increased atherosclerotic plaque deposition in endothelial Nrp2 knockout mice. These findings signpost an atheroprotective role for Nrp2 in ECs.

## Results

### Proteomic profiling of the Nrp2 protein interactome in microvascular ECs

To identify the endothelial protein interactome of Nrp2, microvascular ECs were treated with either non-targeting or Nrp2-specific siRNA duplexes, followed by Nrp2 immunoprecipitation (IP) and label-free quantitative, data-independent acquisition liquid chromatography-tandem mass spectrometry (LFQ-DIA-LC-MS/MS). Putative Nrp2 interactors were determined through statistical analysis comparing the IP scores of non-targeting siRNA-treated samples (Ctrl) to Nrp2 siRNA-treated samples, enabling the isolation of non-specific partner binding. The Spectronaut label-free quantification (LFQ) algorithm was subsequently used for protein quantification with a p-value cutoff of < 0.01 from a total of 3 independent experiments (Figure 1A-B).

**Figure 1:**
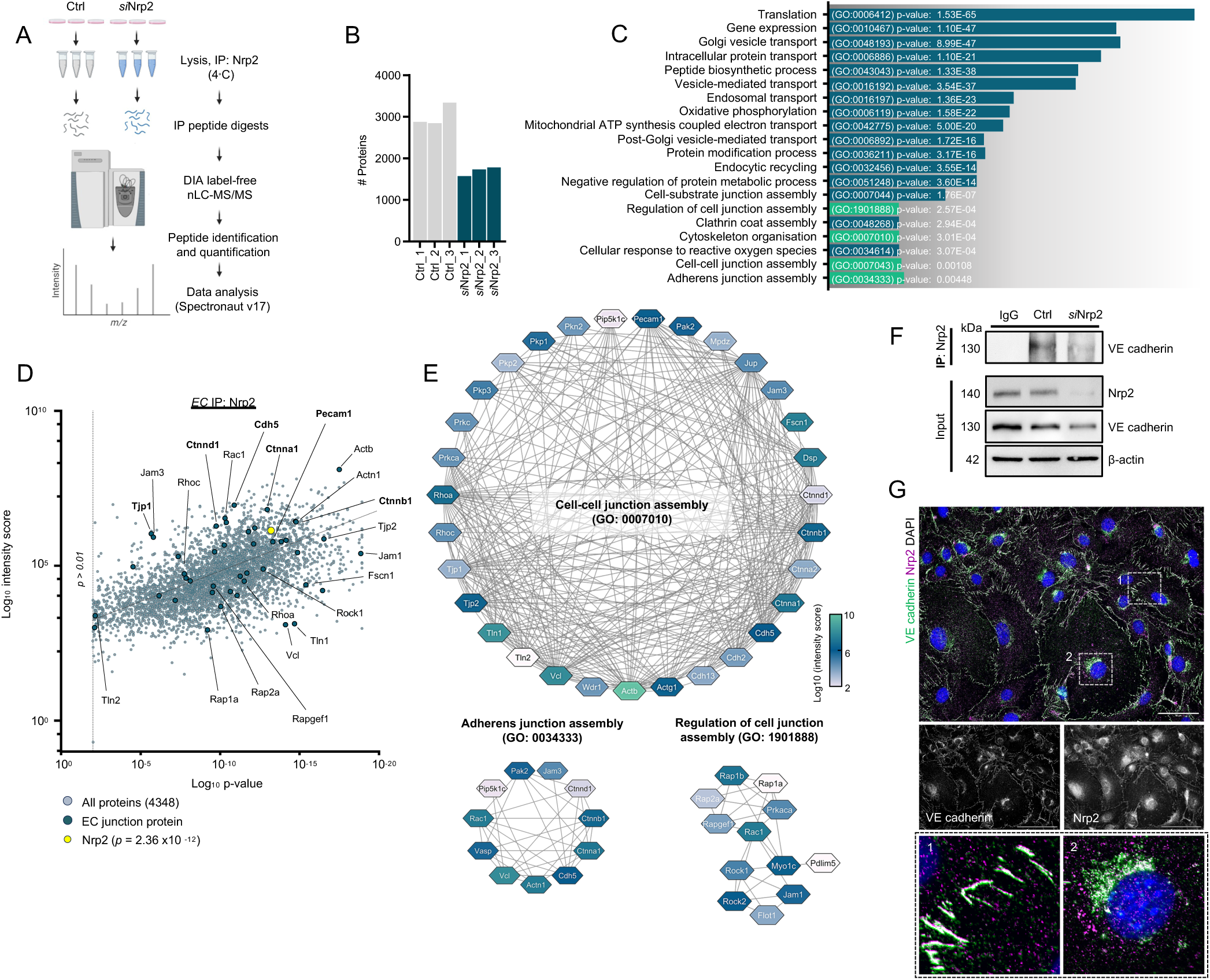
Proteomic profiling of the Nrp2 protein interactome in microvascular ECs. A) LFQ-DIA-LC-MS/MS workflow for proteomics analysis of Nrp2 interactome in microvascular ECs. B) Number of quantified proteins across samples. C) GO enrichment analysis with identifiers and p values showing enriched pathways associated with Nrp2 interactions. D) Scatter plot showing all putative Nrp2 interactions and those associated with junctional assembly (green triangles). E) Protein-protein interactome networks of junctional-associated proteins interacting with Nrp2. Node colour indicates strength of interaction with Nrp2. F) Immunoprecipitation and input Western blots from Ctrl and *si*Nrp2 EC lysates, N = 3 independent experiments. G) Representative confocal microscopy images showing colocalisation between Nrp2 and VE cadherin in confluent fixed ECs. Boxed areas highlight colocalisation both at adherens junctions (1) and in intracellular punctae (2).

Several previously reported binding partners of Nrp2 were identified by our MS analysis, including Vegr1 (*Flt1*), Vegfr2 (*Kdr*), Vegfr3 (*Flt4*), PlexinD1 (*Plxnd1*), α5β1 integrin (*Itga5*, *Itgb1*), Clathrin Heavy Chain-1 (*Cltc*) and Caveolin-1 (*Cav1*). Analysis also revealed candidate interactions with Nrp1 and its endocytic adaptor Gipc1. We proceeded to perform Gene Ontology (GO) enrichment analysis, which confirmed previously published findings that Nrp2 interacts with a number of proteins associated with intracellular receptor traffic, organisation of the actin cytoskeleton and focal adhesion complex assembly, including Rab GTPases 1, 4, 5, 7 and 11, Rac1, RhoA, paxillin (*Pxn*) and focal adhesion kinase (FAK/*Ptk2*). GO enrichment analysis also revealed putative interactions with a high proportion of proteins involved in cell-cell junction assembly (GO:0007043), adherens junction assembly (GO:0034333), and the regulation of cell junction assembly (GO:1901888) including VE cadherin (*Cdh5*), platelet and endothelial cell adhesion molecule-1 (Pecam1), Zonula occludens-1 (ZO-1/*Tjp1*) and the catenin complex (*Ctnna1*, *Ctnnb1*, *Ctnnd1*) (Figure 1C-E).

We proceeded to validate the interaction between Nrp2 and VE cadherin, the principal cell-cell adhesion molecule responsible for maintaining endothelial adherens junctions. After confirming that VE cadherin and Nrp2 form a direct heterological complex in microvascular ECs by IP and Western blotting (Figure 1F), we examined their spatial positioning by immunofluorescence confocal microscopy. We observed a strong colocalisation at cell-cell junctions and in intracellular punctae (Figure 1G), signposting a role for Nrp2 in regulating VE cadherin function.

### Nrp2 depletion stimulates lysosomal-mediated degradation of VE cadherin by destabilising VE cadherin-p120 catenin associations

VE cadherin is the major determinant of contact integrity between ECs, playing an essential role regulating vascular permeability and stability [1], [15]. After verifying our proteomics analysis confirming a direct interaction between Nrp2 and VE cadherin, we explored whether Nrp2 plays a role in regulating junctional assembly and stability. To achieve this, we silenced Nrp2 expression using two Nrp2-specific siRNA constructs and assessed the impact on VE cadherin positioning in confluent ECs subject to either static conditions or orbital shear stress. Unlike those incubated under static conditions, peripherally adhered ECs subject to orbital shear stress displayed partial alignment and elongation, indicative of cultures under laminar flow [16] (Figure 2A). Regardless of whether subject to static conditions or laminar flow, EC depleted for Nrp2 exhibited a significant intracellular accumulation of VE cadherin compared to their control counterparts, suggesting a role for Nrp2 in maintaining junctional stability and VE cadherin surface availability (Figure 2B-D). Indeed, *si*Nrp2 ECs grown under laminar flow exhibited fewer VE cadherin^+^ and p120-catenin^+^ continuous junctions than Ctrl ECs (Figure B, E). *si*Nrp2 ECs grown under static conditions also exhibited significantly reduced junctional VE cadherin expression at discontinuous adherens junctions compared to Ctrl ECs, alongside a reduction in the number of adjoining linear actin stress fibres and an abundant increase in the incidence of cortical actin (Figure 2B, Suppl. Figure 1A-D).

**Figure 2:**
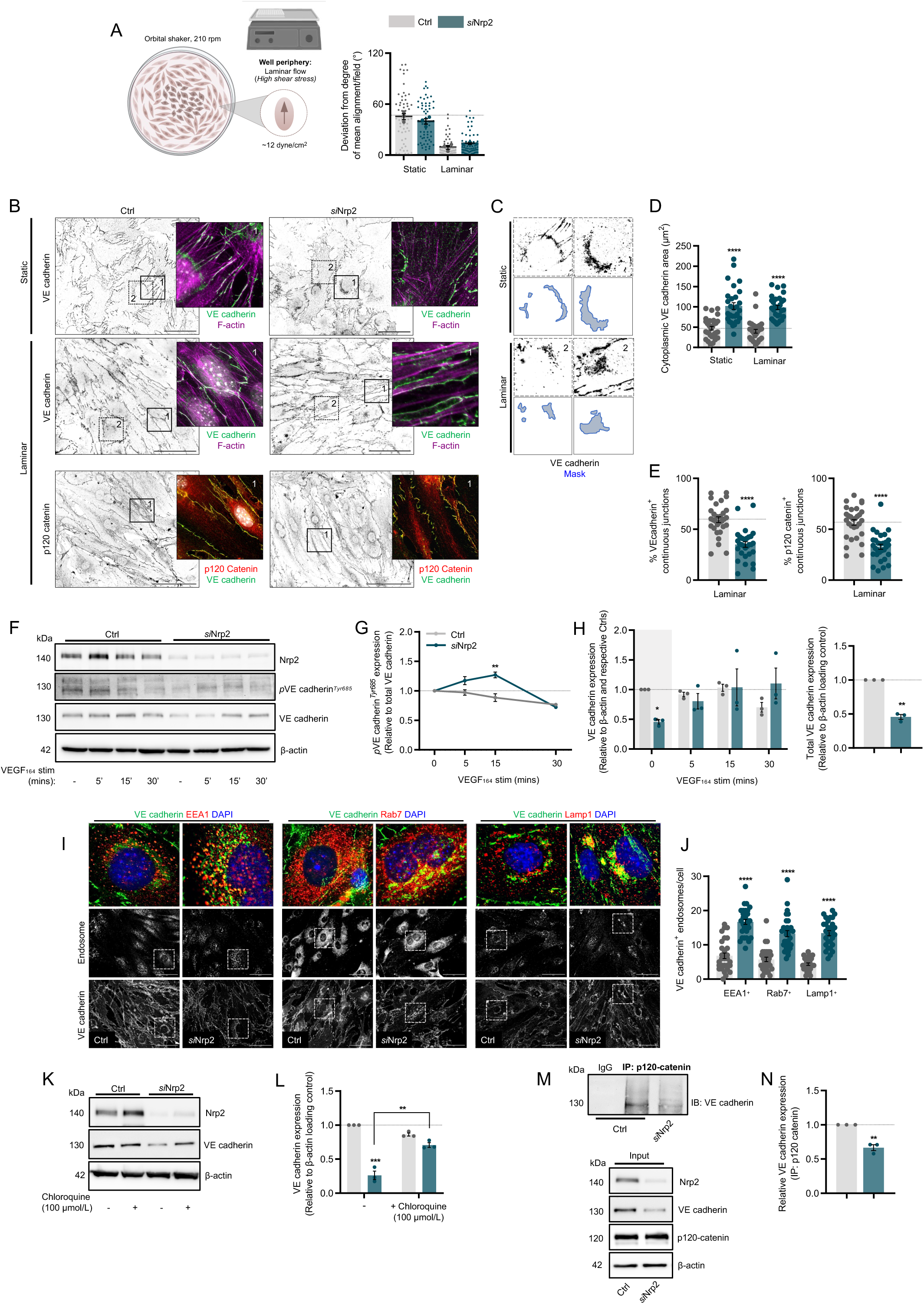
Nrp2 depletion stimulates lysosomal-mediated degradation of VE cadherin by destabilising VE-cadherin-p120-catenin associations. A) (Left panel) Schematic showing orbital shaker method to induce laminar flow. (Right panel) Quantification of Ctrl and *si*Nrp2 EC alignment under static conditions or following incubation under laminar flow, N = 3 independent experiments, (n ≥ 60). B) Representative confocal microscopy images showing VE cadherin/F-actin (top panels) and VE cadherin/p120 catenin (bottom panels) in confluent fixed Ctrl and *si*Nrp2 ECs subject to either static conditions or laminar flow. C) High magnification images of highlighted areas shown in B (top panels) showing VE cadherin accumulation in the perinuclear region. D) Quantification of intracellular VE cadherin area (µm^2^) in Ctrl and *si*Nrp2 ECs, N = 3 independent experiments, (n ≥ 30). E) Quantification of % VE cadherin/p120 catenin^+^ continuous junctions/cell in Ctrl and *si*Nrp2 ECs subject to laminar flow, N = 3 independent experiments, (n ≥ 30). F) Representative Western blotting showing *p*VE cadherin^Tyr685^, total VE cadherin and β-actin loading control in Ctrl and siNrp2 ECs with VEGF_164_ stimulation. G) Quantification of *p*VE cadherin^Tyr685^ expression relative to total VE cadherin expression and to unstimulated conditions, N = 3 independent experiments. H) (Left panel) Quantification of total VE cadherin expression relative to loading control and respective control timepoints, N = 3 independent experiments. (Right panel) Quantification of total VE cadherin expression relative to loading control under unstimulated conditions, N = 3 independent experiments. I) Representative confocal microscopy images showing colocalisation between VE cadherin and EEA1, Rab7, and Lamp1 in fixed, confluent Ctrl and *si*Nrp2 ECs. J) Quantification of VE cadherin^+^ EEA1, Rab7 and Lamp1 endosomes/cell under unstimulated conditions, N = 3 independent experiments, (n ≥ 30). K) Representative Western blotting showing total VE cadherin and β-actin loading control in Ctrl and *si*Nrp2 ECs ± 100 µmol/L chloroquine (4 hrs). L) Quantification associated with K), N = 3 independent experiments. M) (Top panel) Immunoprecipitation from Ctrl and *si*Nrp2 EC lysates of p120 catenin followed by SDS-PAGE and Western blotting with an anti-VE cadherin antibody. (Bottom panel) Total cell lysate input showing Nrp2 depletion, VE cadherin, p120 catenin expression, and loading control. N) Quantification of relative VE cadherin expression (IP: 120 catenin), N = 3 independent experiments. Asterixis indicate significance.

VE cadherin phosphorylation at its *Tyr*^685^ residue by Src kinase is known to induce rapid endocytosis and adherens junction disassembly [17]. To examine the effect of Nrp2 depletion on this signalling cascade, we stimulated Ctrl and *si*Nrp2 depleted ECs with Vegf-A_164_ (the mouse-specific isoform of Vegf-A_165_) for up to 30 minutes before quantifying phosphorylated and total VE cadherin expression by Western blotting. After 15 minutes Vegf-A_164_ stimulation, we observed a significant increase in phosphorylated VE cadherin *^Tyr^*^685^ expression (relative to total VE cadherin expression) in our *si*Nrp2 ECs (Figure 2F-G). Nrp2 silencing was also found to induce a significant reduction in total VE cadherin expression under unstimulated conditions (Figure 2F, H). To investigate the subcellular localisation of VE cadherin under these conditions, we subsequently performed confocal immunofluorescence microscopy of fixed ECs labelled with endogenous VE cadherin and markers of early endosomes (EEA1^+^), late endosomes (Rab7^+^) and lysosomes (Lamp1^+^). VE cadherin was found to preferentially accumulate in each of these intracellular compartments upon Nrp2 depletion (Figure 2I-J), suggesting that its expression protects against the rapid endocytosis and lysosomal degradation of VE cadherin in ECs. To verify this hypothesis, we pre-treated unstimulated Ctrl and *si*Nrp2 ECs with the well-characterised lysosomal inhibitor chloroquine in an attempt to rescue total VE cadherin expression. Chloroquine incubation for 4 hours at 100 µmol/L dramatically inhibited the downregulation of VE cadherin observed in cells depleted for Nrp2 (Figure 2K-L), indicating that following Nrp2 depletion, VE cadherin is processed intracellularly and transported for degradation.

p120-catenin has previously been reported to stabilise VE cadherin at cell-cell contacts, inhibiting its CME following *Tyr*^685^ phosphorylation [8], [9]. Nrp1 was also revealed to promote the association between VE cadherin and p120-catenin in human umbilical vein ECs (HUVECs) to preserve junctional integrity [10]. We therefore considered whether Nrp2 would share a similar function. Indeed, biochemical immunoprecipitation studies showed a reduced interaction between VE cadherin and p120-catenin in the absence of Nrp2, despite no changes in total p120-catenin expression (Figure 2M-N). Taken together, these data suggest that Nrp2 maintains junctional VE cadherin expression by promoting its association with p120-catenin. Upon loss of Nrp2, VE cadherin is phosphorylated, endocytosed, and intracellularly processed following a reduced association with p120-catenin and impaired actin cytoskeleton remodelling.

### Nrp2 promotes junctional integrity during neovascularisation of the postnatal retina

To investigate the impact of Nrp2 depletion *in vivo*, we utilised previously generated Nrp2*^flfl^*.Pdgfb-iCreER^T2^ (Nrp2*^flfl^*.*EC^KO^*) mice [14], [18], [19] and examined VE cadherin distribution and vessel integrity in the developing retina at postnatal day 6 (Figure 3A). First, whole-mount immunostaining in control littermate animals revealed Nrp2 expression to be concentrated in regions enriched for VE cadherin, in agreement with our earlier findings, and those reported by Fantin *et al*., when describing immunolocalization of Nrp1 [20] (Figure 3B-C). We then proceeded to compare the distribution of VE cadherin at the vascular front and vascular interior of retinas harvested from control (Nrp2*^flfl^*) and Nrp2*^flfl^*.*EC^KO^* mice. Whilst VE cadherin positioning in Nrp2*^flfl^*retinas appeared sharp and continuous, clearly marking junctional boundaries, in Nrp2*^flfl^*.*EC^KO^* retinas, VE cadherin staining was observed as diffuse and discontinuous, particularly in newly forming sprouts (Figure 3D-E). Indeed, quantification of cytoplasmic VE cadherin intensity in Nrp2*^flfl^*.*EC^KO^* vasculature was revealed to be significantly higher than in littermate control vasculature, concomitant with a significantly higher incidence of junctional gaps indicative of reduced vessel integrity (Figure 3F-G). Manual classification of VE cadherin patterning (active to inactive) supported these findings, with Nrp2*^flfl^*.*EC^KO^*vasculature displaying an increased incidence of serrated and discontinuous junctions compared to control vasculature (Suppl. Figure 2A).

**Figure 3:**
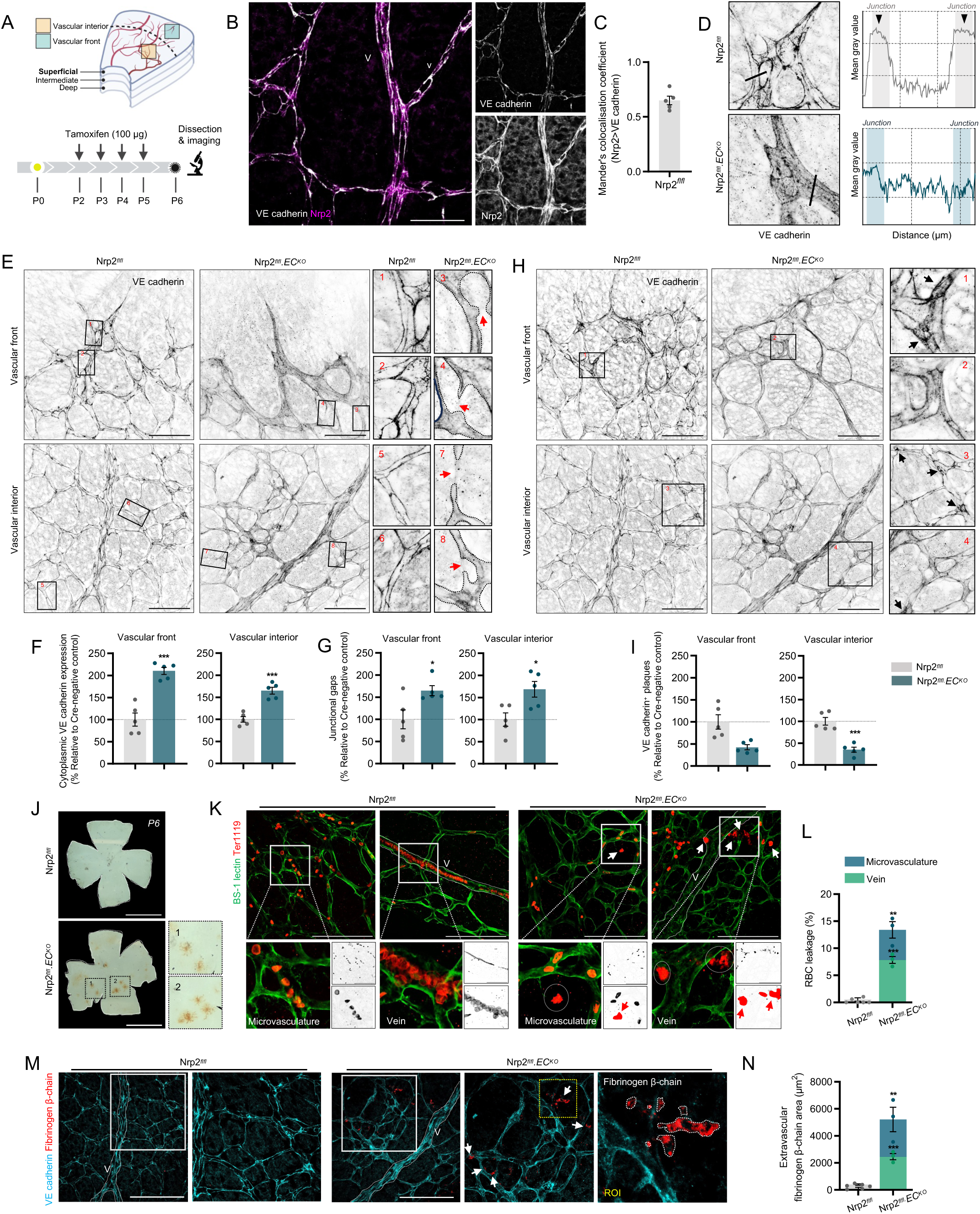
Nrp2 promotes junctional stability during neovascularisation of the postnatal retina. A) Inducible, endothelial-specific deletion of Nrp2 was achieved by crossing Nrp2*^flfl^*mice with Pdgfb-iCreER^T2^ (Nrp2*^flfl^*.*EC^KO^*) mice. Activation of Pdgfb-iCreER^T2^ was achieved by subcutaneous injection of tamoxifen on P2 and P3, followed by intraperitoneal (IP) injections on P4 and P5. Retinas were then harvested at P6. B) Confocal microscopy image showing colocalisation of Nrp2 in veins (V) and venules (v) with VE cadherin. C) Quantification of colocalised Nrp2 > VE cadherin pixels, (n ≥ 5 retinas). D) Representative confocal microscopy images showing P6 retinal vasculature labelled with VE cadherin at the vascular front and in the vascular interior with corresponding intensity maps showing expression of VE cadherin across the lines shown in the magnified selections. E) Representative confocal microscopy images showing P6 retinal vasculature labelled with VE cadherin at the vascular front and in the vascular interior. Red arrows indicate junctional gaps. F) Quantification of cytoplasmic VE cadherin expression (shown as % relative to Cre-negative control data), n = 5 retinas. G) Quantification of the number of junctional gaps/field (VE cadherin^-^) (shown as % relative to Cre-negative control data), n = 5 retinas (4x ROIs/retina). H) Representative confocal microscopy images showing P6 retinal vasculature labelled with VE cadherin at the vascular front and in the vascular interior (highlighted areas show VE cadherin plaques). I) Quantification of the number of VE cadherin^+^ plaques/field (shown as % relative to Cre-negative control data), n = 5 retinas, (4x ROIs/retina). J) Representative brightfield images of Nrp2*^flfl^* and Nrp2*^flfl^.EC^KO^*P6 dissected retinas showing minor blood haemorrhages in Nrp2*^flfl^.EC^KO^* P6 dissected retinas only. K) Representative confocal microscopy images showing Ter-119^+^ red blood cell (RBC) leakage from BS-1 lectin labelled vasculature. L) Quantification of % RBC leakage from Nrp2*^flfl^* and Nrp2*^flfl^.EC^KO^* P6 retinas, n = 3 retinas (5x ROIs/retina). M) Representative confocal microscopy images showing fibrinogen β-chain^+^ low molecular weight leakages from VE cadherin labelled vasculature. N) Quantification of fibrinogen β-chain^+^ leakage area (µm^2^) from Nrp2*^flfl^* and Nrp2*^flfl^.EC^KO^* P6 retinas, n = 3 retinas (5x ROIs/retina). Asterixis indicate statistical significance.

Polarised and iterative assembly of junction-associated intermittent lamellipodia (JAIL) has previously been shown to be associated with increased vessel integrity and polarised tip cell sprouting *in vivo* [21], [22]. The relative decrease in VE cadherin concentration at EC junctions during cell elongation and sprouting is understood to stimulate the assembly of JAIL plaques, promoting the formation of new VE cadherin adhesion sites to maintain the junctional barrier and limit vascular leakage. In line with a significant increase in junctional gaps, Nrp2*^flfl^*.*EC^KO^* vasculature was also found to exhibit significantly fewer VE cadherin^+^ plaque-like assemblies between adjoining ECs compared to retinas extracted from respective littermate control mice (Figure 3H-I). These data indicate that Nrp2 promotes the assembly of nascent VE cadherin^+^ boundaries.

We proceeded to directly assess the impact of Nrp2 deletion on EC monolayer integrity by immunostaining for Ter-119, a marker of red-blood cells (RBCs) [23]. At the outset, we had, by eye, observed occasional minor haemorrhages in the retinas of Nrp2*^flfl^*.*EC^KO^* animals, which were absent in retinas extracted from littermate control animals (Figure 3J). Indeed, immunostaining revealed a modest yet significant increase in RBC leakage from the retinal microvasculature following Nrp2 deletion (Figure 3K-L). To further verify the permeability of Nrp2*^flfl^*.*EC^KO^* retinal vasculature, low-molecular weight leakage of the endogenous plasma protein fibrinogen was quantified. In contrast to control retinal vasculature, extravascular fibrinogen was detected sporadically in Nrp2*^flfl^*.*EC^KO^*mice, particularly around the retinal microvasculature (Figure 3M-N). Collectively, these data suggest that Nrp2 promotes junctional integrity *in vivo* by regulating VE cadherin distribution in newly developing vascular beds.

### ***si***Nrp2 ECs exhibit a pro-inflammatory phenotype

Loss of VE cadherin-mediated junctional integrity and increased vascular permeability is a hallmark of endothelial activation and dysfunction, leading to vascular inflammation and atherogenesis [10], [24], [25]. We therefore analysed the differential expression of genes known to play a role in inflammation and atherosclerosis in Ctrl vs *si*Nrp2 ECs cultured under static conditions by standard RNA sequencing (RNA-seq) (Figure 4A-E, Suppl. Figure 3A). Gene ontology (GO) enrichment analysis using KEGG and Reactome annotation databases identified that a high proportion of upregulated genes had functions associated with cell responses to cytokine stimulus during the immune response, MAPK signalling, and leukocyte adhesion. Conversely, Nrp2 depletion resulted in the downregulation of genes with functions associated with extracellular matrix organisation, ion transport, lipid metabolism and cell-cell junction assembly (Figure 4F). Uniform Manifold Approximation and Projection (UMAP) clustering of genes associated with the inflammatory response confirmed our gene enrichment pathway analysis (Suppl. Figure 3B).

**Figure 4:**
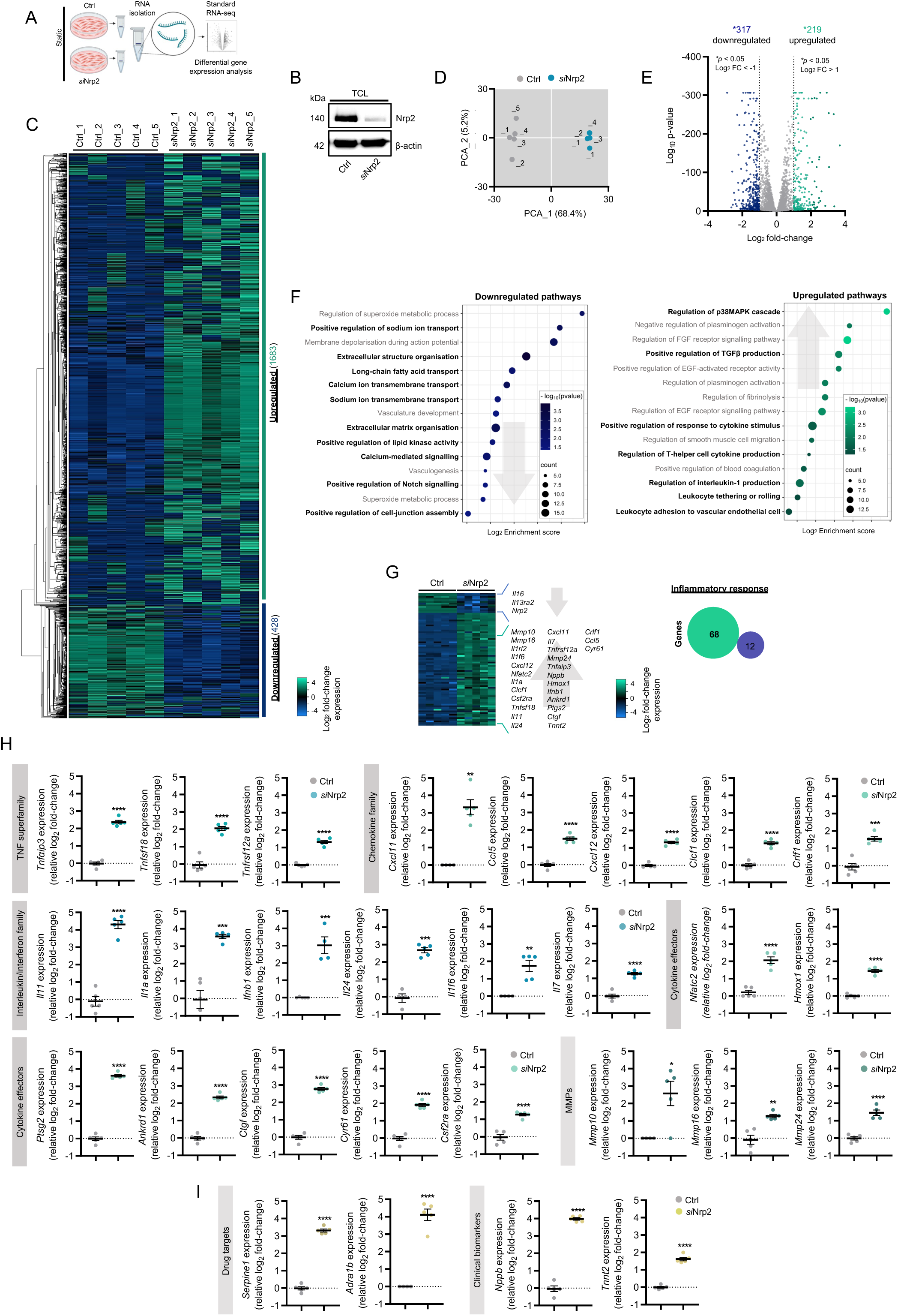
siNrp2 ECs exhibit a pro-inflammatory phenotype. A) Schematic showing RNA-seq workflow. B) Total cell lysate input confirming Nrp2 depletion from SDS-PAGE and Western blotting. C) PCA plot showing two component separation between Ctrl and *si*Nrp2 samples, N = 5 independent experiments. D) Heatmap of transcriptional changes between Ctrl and *si*Nrp2 samples. E) Volcano plot of RNA-seq transcriptomic data displaying mean significantly differentially expressed genes in *si*Nrp2 ECs relative to Ctrl ECs cultured under static conditions, N = 5 independent experiments. F) GO enrichment analysis of pathways differentially modulated in *si*Nrp2 ECs relative to Ctrl ECs cultured under static conditions. G) Heatmap and Venn diagram showing differential gene expression changes in the inflammatory response between Ctrl and *si*Nrp2 samples, N = 5 independent experiments. H) Barplots showing upregulated genes associated with the inflammatory response between Ctrl and *si*Nrp2 samples, N = 5 independent experiments. I) Barplots showing upregulation of known drug targets and clinical biomarkers of cardiovascular disease.

Of the 2111 differentially expressed genes (log_2_ fold-change > 1, < 1), 317 were significantly downregulated, including Nrp2 (Suppl. Figure 3C). We also observed a significant downregulation in the expression of interleukin-13 receptor (*Il13ra2*), which is known to exclusively bind IL13, in addition to a significant downregulation in interleukin-16 (*Il16*) expression (Suppl. Figure 3D). Both *Il13r* (when bound to *Il13*) and *Il16* are known to play a protective role during atherosclerosis progression [26], [27]. No gene expression changes were observed for Nrp1, as previously reported [14] (Suppl. Figure 3E).

Of the 219 significantly upregulated genes, Nrp2 depletion induced the expression of mRNAs encoding a number of pro-inflammatory markers (a total of 68 genes), including members of the TNF superfamily (*Tnfαip3, Tnfsf18*, *Tnfrsf12a*), chemokine family (*Ccl5*, *Cxcl11*, *Cxcl12, Clcf1, Crlf1*), and interleukin/interferon families (*Il1a*, *Il1f6*, *Il1rl2*, *Il7*, *Il11*, *Il24, Ifnb1*) [25], [26], [28], [29], [30], [31] (Figure 4G). Ankyrin repeat domain-containing protein 1 (*Ankrd1*), which is canonically induced following IL1 and TNFα-stimulation, both connective tissue growth factor (*Ctgf*) and cysteine-rich angiogenic inducer-61 (*Cyr61*) [32], which are highly expressed in atherosclerotic human arteries, and nuclear factor of activated T-cell 2 (*Nfatc2*), a pro-inflammatory transcription factor induced by IL11 signalling in ECs [31], were all also significantly upregulated (Figure 4H). Whilst we observed no significant changes to the pro-inflammatory effector TGFβ, which is commonly associated with endothelial dysfunction, *αV β6* integrin expression, which activates TGFβ1 signalling during the inflammatory response [33], [34], was found to significantly increase (Suppl. Figure 3F).

In addition to changes in the expression profiles of inflammatory cytokines, RNA-Seq analysis identified the upregulation of matrix metalloproteases 10, 16 and 24 (*Mmp10*, *Mmp16*, *Mmp24*) (Figure 4H), all previously shown to contribute to the pathogenesis of cardiovascular diseases by promoting EC dysfunction and immune cell invasion. Indeed, *Mmp10*, *Mmp16*, *Mmp24* were all reported to be upregulated in atherosclerotic plaques and human cardiac ECs during myocardial infarction [35], [36]. Nrp2 depleted ECs were also found to exhibit elevated plasminogen activating inhibitor-1 (PAI-1/*Serpine-1*) and α-1b adrenergic receptor (*Adrab1*) gene expression, both known drug targets of atherosclerosis progression. Finally, Nrp2 silencing induced the expression of natriuretic peptide B (*Nppb*) and troponin-T (*Tnnt2*) mRNAs, both of which are used clinically as predictive biomarkers for cardiovascular ischemia [37], [38], [39], [40] (Figure 4I).

### Glycolysis is upregulated upon Nrp2 silencing

Despite this transcriptional shift to a pro-inflammatory phenotype, we did not observe any gross changes in the gene expression of common flow-sensitive transcription factors or transcriptional co-activators such as Krüppel-like factor 2 and 4 (*Klf2*, *Klf4)*, *Nf-κB*, *Hif1α*, *Yap*, *Taz* or *Sox13*, typically reported as being altered in states of disturbed flow [10], [41]. RNA-Seq analysis did detect an upregulation of multiple genes regulating glycolysis however, including hexokinase-2 (*Hk2)* [42], the first enzyme involved during glucose metabolism, in addition to aldolase (*Aldoa*), enolase (*Eno1*) and lactate dehydrogenase A (*Ldha*) (Figure 5A-B). Instigating a re-inspection of our proteomics dataset, we were surprised to find that Nrp2 exhibited robust interactions with all enzymes participating in the glycolytic cascade. Specifically, interactions with pyruvate kinase (Pkm), glyceraldehyde 3-phosphate dehydrogenase (Gapdh), Eno1, Aldoa, and Ldha were all detected within the top 20 proteins with the greatest interaction (LFQ) score. Unsurprisingly, pathway enrichment analysis of these proteins clearly revealed a strong association with glucose metabolism (GO:0051156) (Figure 5C). *si*Nrp2 ECs were also found to exhibit significantly elevated gene expression of fibroblast growth factors (*Fgfs*) 2, *5*, *9* and *16*, all reported to induce glycolysis under stress (Suppl. Figure 4A).

**Figure 5:**
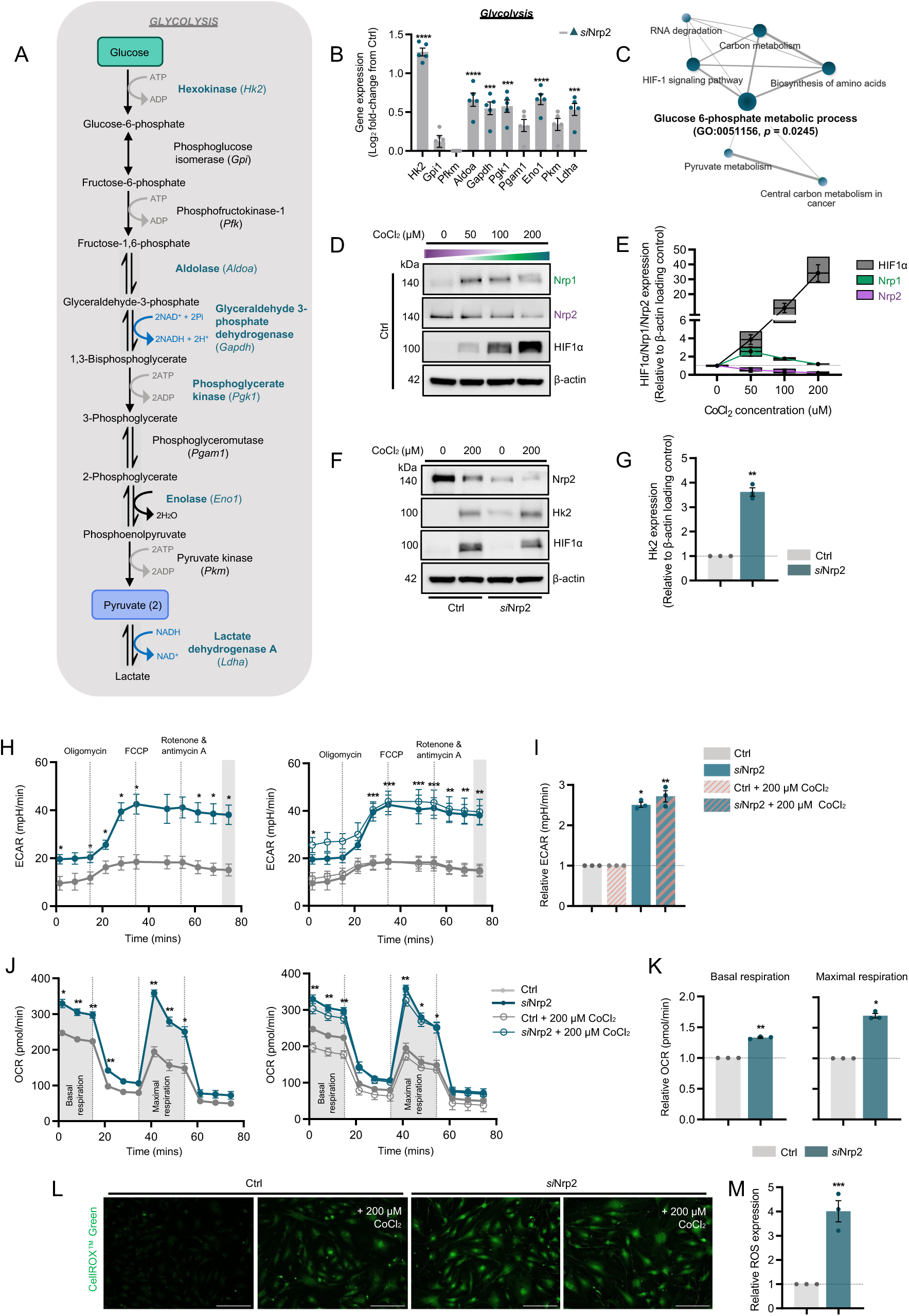
Glycolysis is upregulated upon Nrp2 silencing. A) Pathway schematic of glucose metabolism. Genes upregulated following Nrp2 depletion labelled in teal. B) Barplots showing transcriptional changes in genes regulating glycolysis and oxidative stress. C) KEGG pathway network of proteins interacting with Nrp2 with the highest LFQ score (top 20). D) Western blotting from total cell lysate of Nrp1, Nrp2 and HIF1α following CoCl_2_ treatment in Ctrl ECs. E) Quantification of Nrp1, Nrp2 and HIF1α expression following CoCl_2_ treatment in Ctrl ECs. F) Western blotting from total cell lysate of Nrp2 and Hk2 following CoCl_2_ treatment in Ctrl and *si*Nrp2 ECs. G) Quantification of Hk2 expression under basal conditions relative to loading control, N = 3 independent experiments. H) Seahorse kinetics of mean ECAR (mpH/min) measurements in Ctrl and *si*Nrp2 ECs under basal conditions (left panel) or following CoCl_2_ treatment (right panel), N = 3 independent experiments. I) Seahorse analysis barplot of mean ECAR (mpH/min) measurement (final reading), N = 3 independent experiments. J) Seahorse kinetics of mean oxygen consumption rate (OCR) (*p*mol/min) measurements in Ctrl and *si*Nrp2 ECs under basal conditions (left panel) or following CoCl_2_ treatment (right panel), N = 3 independent experiments. K) Seahorse analysis of mean relative basal respiration and maximal respiration rates in Ctrl and *si*Nrp2 ECs under basal conditions, N = 3 independent experiments. L) Representative confocal microscopy images showing ROS expression in Ctrl and *si*Nrp2 ECs under normoxia and following CoCl_2_ treatment. M) Quantification of ROS expression, N = 3 independent experiments. Asterixis indicate statistical significance.

During atherogenesis, lipid accumulation and inflammation promotes the transition from oxidative metabolism to glycolysis and a state of oxidative stress [43], [44]. As hypoxia is known to enhance glycolysis during atherosclerosis [43], [45], we proceeded to examine glycolytic capacity in ECs depleted for Nrp2 treated with or without the hypoxia mimetic agent CoCl_2_ to pathophysiologically induce HIF1α signalling. As reported previously in glioblastoma and melanoma cell lines [46], a state of hypoxia-driven signalling inhibited Nrp2 expression levels dramatically. In contrast, CoCl2 treatment increased Nrp1 expression up to a concentration of 100 µM (Figure 5D-E). Mimicking the upregulation of *Hk2* gene expression following Nrp2 silencing, Hk2 protein levels in CoCl_2_-treated Ctrl lysates and in *si*Nrp2 lysates were found to be elevated (Figure 5F-G). Given Nrp2 depletion increased gene and protein expression of essential glycolytic enzymes, we proceeded to measure glycolytic output directly using Seahorse analysis, under both basal conditions and in a state of CoCl_2_-induced pseudo-hypoxic stimulation. *si*Nrp2 ECs were found to exhibit significantly increased glycolytic metabolism under basal conditions, assessed by measuring extracellular proton flux (extracellular acidification rate (ECAR)). Despite effectively inducing HIF1α signalling and Hk2 expression, CoCl_2_ treatment did not exert any additional effect (Figure 5H-I). Based on these results, we postulated that the elevated rate of glycolysis exhibited by *si*Nrp2 ECs may result from a supressed rate of aerobic mitochondrial respiration in order to sustain ATP generation. Surprisingly however, Nrp2 depleted ECs were also found to exhibit significantly increased basal and maximal respiration capacity compared to Ctrl ECs (Figure 5J-K), suggestive that Nrp2 participates in modulating aerobic EC metabolism in addition to protecting against glycolytic stress. In support of this, we observed significantly elevated expression of reactive oxygen species (ROS) in *si*Nrp2 ECs (Figure 5L-M), the production of which is induced by increased rates of respiration and the pro-inflammatory pathways activated during cardiovascular disease progression [47].

### Nrp2 protects against EC activation, immune-cell attachment and atherosclerotic plaque deposition

We proceeded to investigate the functional effects of the observed pro-inflammatory transcriptional changes by assessing whether Nrp2 depletion would upregulate the expression of immunoglobulin-like adhesion molecules Vcam1 and Icam1. Both Vcam1 and Icam1, alongside the lesser studied Icam2, are known to be expressed by activated ECs following cytokine release and or in response to disturbed flow. Their expression subsequently facilitates monocyte attachment and extravasation to promote inflammation and plaque deposition during atherosclerosis progression [41], [48], [49]. As Nrp1*^flfl^*.*EC^KO^*mouse aortas were reported to exhibit increased Vcam1 expression compared to control littermate animals [10], we first examined whether an endothelial-specific deletion of Nrp2 would elicit a similar effect. Indeed, immunostaining in the descending aortas of 8-week-old Nrp2*^flfl^*.*EC^KO^*mice revealed increased Vcam1 expression at the apical surface compared to their control counterparts (Figure 6A-C).

**Figure 6:**
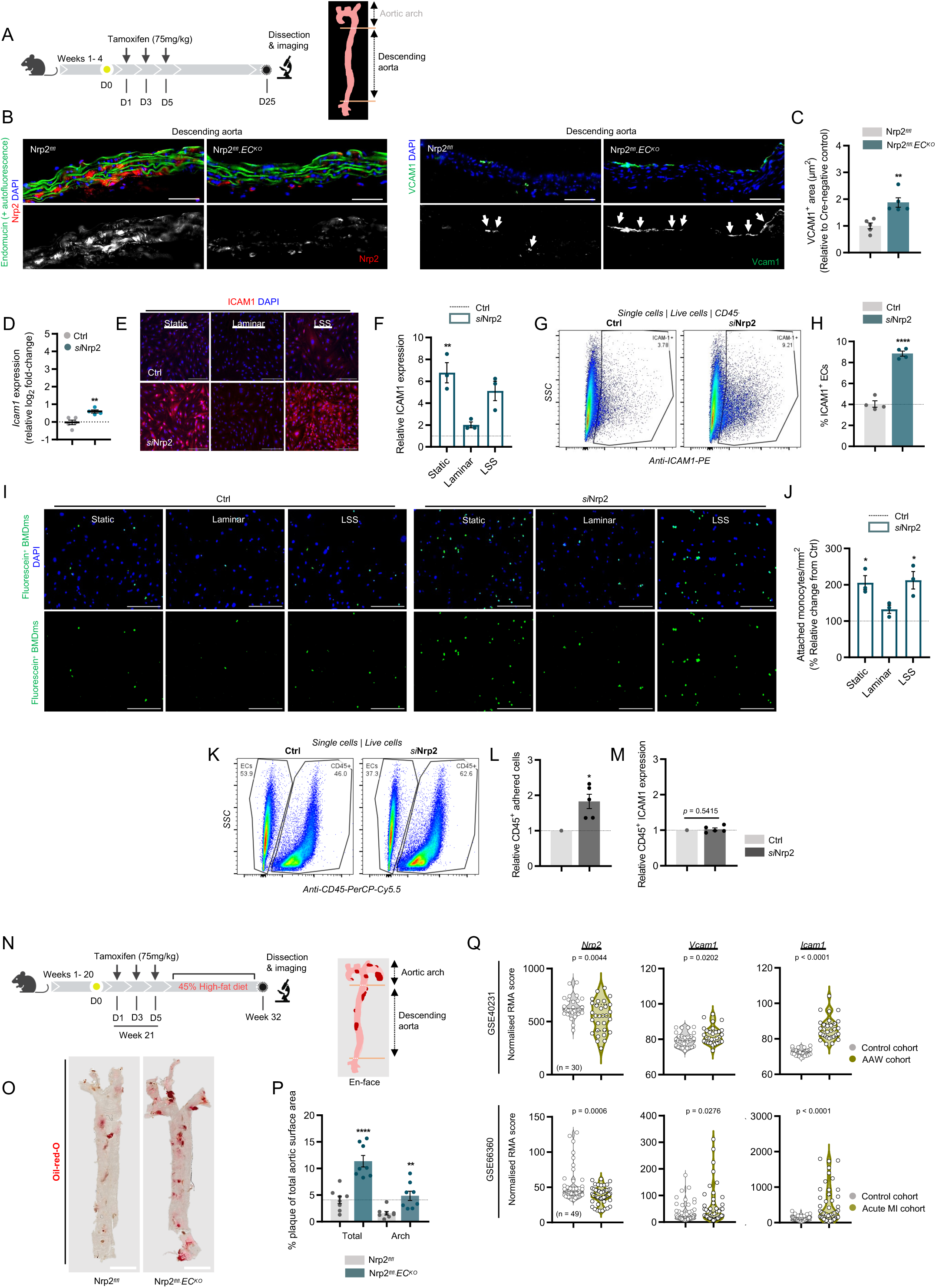
Nrp2 protects against EC activation and immune-cell attachment, but not atherosclerosis development. A) Experimental schematic showing tamoxifen-induced gene depletion regime, and diagram of ascending (arch) and descending portions of the mouse aorta. B) (Left panels) Representative confocal microscopy images confirming Nrp2 knockdown in the descending aortas of Nrp2*^flfl^.EC^KO^*mice. (Right panels) Representative confocal microscopy images showing increased VCAM1 expression in the descending aortas of Nrp2*^flfl^.EC^KO^* mice. C) Quantification of aortic VCAM1^+^ area (shown as relative to Cre-negative control data), n = 5 animals. D) Quantification of *Icam1* gene expression between Ctrl and *si*Nrp2 samples, N = 5 independent experiments. E) Representative confocal microscopy images showing ICAM1 expression in Ctrl and *si*Nrp2 ECs cultured under static, laminar, or low shear stress conditions. F) Quantification of relative ICAM1 expression in Ctrl and *si*Nrp2 ECs cultured under static, laminar, or low shear stress conditions, N = 3 independent experiments, (n ≥ 60). G-H) Representative flow cytometry plots and respective quantification of % Icam1^+^ ECs following a 1 hr co-culture with BMDms cultured under static conditions, N = 4 independent experiments. I) Representative confocal microscopy images showing Ctrl and *si*Nrp2 EC-adhered fluorescein-labelled BMDms cultured under static, laminar, or low shear stress conditions. J) Quantification of EC-adhered BMDms/mm^2^ (shown as % relative to Ctrl), N = 3 (12x ROIs/independent experiment). K-L) Representative flow cytometry plots and respective quantification of relative number of adhered CD45^+^ BMDms following a static co-culture with Ctrl and *si*Nrp2 ECs, N = 5 independent experiments. M) Quantification of relative number of ICAM1^+^ BMDms following a static co-culture with Ctrl and *si*Nrp2 ECs. N) Experimental schematic showing tamoxifen-induced gene depletion and high-fat diet regime. O) Brightfield images of en-face Oil-red-O-stained aortas dissected from Nrp2*^flfl^*and Nrp2*^flfl^.EC^KO^* animals at week 20. P) % Relative quantification of total Oil-red-O^+^ plaque area, n = 8. Q) Gene expression analysis from GSE40231 and GSE66360 microarray datasets. Asterixis indicate statistical significance.

Whilst this observed upregulation in aortic Vcam1 expression did not correspond with elevated *Vcam1* mRNA levels from cultured *si*Nrp2 EC samples, RNA-Seq analysis did reveal a modest yet significant increase in *Icam1* gene expression (Figure 6D). We observed similar increases in Icam1 protein expression in *si*Nrp2 ECs subject to static or low shear stress (LSS) environments, to simulate disturbed flow (Figure 6E-F, Suppl. Figure 5A). Moreover, *si*Nrp2 ECs subject to a 1-hour co-culture with bone-marrow derived monocytes (BMDms) also exhibited upregulated Icam1 expression (Figure 6G-H). These data suggest that Nrp2 silencing promotes adhesion receptor expression in environments where laminar flow is disrupted or absent.

We proceeded to confirm this hypothesis by determining whether Nrp2 depletion would increase monocyte-endothelium attachment. In addition to significantly increasing *Icam1* expression, Nrp2 silencing was also found to upregulate several genes with roles promoting monocyte and neutrophil adhesion, rolling and transmigration, including *S100a8, Edn1,* and *Selplg* [50], [51], [52]. Furthermore, Nrp2 depletion significantly reduced *Nos2* expression, which has been shown to inhibit monocyte attachment via its production of NO under normoxia (Suppl. Figure 5B). Consistent with our previous findings, and these observed gene changes, significantly more BMDms were found to adhere to Nrp2 silenced ECs subject to LSS or under static conditions, compared to Ctrl ECs (Figure 6I-J). This observed increase in monocyte attachment was subsequently validated by flow cytometry and was found not to result from any changes to monocytic Icam1 expression (Figure 6K-M). Together, our findings indicate that Nrp2 expression protects against endothelial activation and elevated Vcam1/Icam1 expression, moderating monocyte attachment and their subsequent infiltration to reduce tissue inflammation.

During atherosclerosis progression, endothelial activation and dysfunction occurs early, contributing to a chronic inflammatory environment to promote atheroma formation. To determine whether Nrp2 has atheroprotective functions, we examined atherosclerotic plaque development in aortas of 20-week-old control and Nrp2*^flfl^*.*EC^KO^* animals fed a 45% high-fat diet for a successive 12 weeks (Figure 6N). en-face Oil-red-O staining of Nrp2*^flfl^*.*EC^KO^* aortas revealed an increase in atherosclerotic lesion area compared to aortas isolated from littermate control animals (Figure 6O-P) despite no gross changes in overall weight.

To validate our results, we identified published microarray datasets *GSE40231* and *GSE66360* from the GEO database, comparing gene expression profiles from healthy and disease patient cohorts. The *GSE40231* dataset compares non-atherosclerotic arterial wall tissue (distal mammary artery) (control) with atherosclerotic arterial wall (AAW) tissue isolated from 30 patients during coronary artery bypass grafting surgery [53]. The *GSE66360* dataset compares gene profiles from circulating endothelial cells isolated from patients experiencing acute myocardial infarction (n = 49) with a healthy cohort (n = 50) [54]. Consistent with published literature, gene expression levels of both *Vcam1* and *Icam1* were significantly elevated in disease samples compared to control samples. In contrast, disease samples exhibited significantly reduced *Nrp2* expression (Figure 6Q), corroborating the evidence presented in this study that targeting Nrp2 expression promotes endothelial dysfunction and atherogenesis.

## Discussion

Neuropilin receptors, regardless of their traditional interactions with VEGF and Sema3, have been shown to play an essential role in blood vessel development [10], [14], [20], [55], [56], [57], [58]. The precise mechanisms by which they coordinate dynamic rearrangements of adhesive contacts to aid in the formation of a functional vascular network, however, remains poorly understood. The sum of this work establishes a novel role for Nrp2 in promoting junctional integrity and protecting against the onset of atherogenesis. Nrp2 silencing was found to destabilise the VE cadherin-p120-catenin complex, stimulating the rapid endocytosis and degradation of VE cadherin. As a consequence, junctional availability of VE cadherin is reduced, increasing vascular leakage *in vivo.* RNA-Seq analysis identified a pro-inflammatory transcriptional shift in *si*Nrp2 ECs to induce EC activation, monocyte attachment and, as a consequence, increased aortic plaque deposition.

The role of Nrp2 in microvascular ECs has been shown in recent years to closely follow that of its better studied structural ortholog, Nrp1. Indeed, Nrp2, like Nrp1, is now understood to promote α5β1 integrin-mediated adhesion to fibronectin during polarised EC migration and sprouting [11], [14], [18], [58]. In a manner similar to that reported by Bosseboeuf *et al*., [10] we now demonstrate that Nrp2 promotes the biochemical association between VE cadherin and p120-catenin, preserving its expression at cell-cell junctions to modulate junctional integrity and vascular leakage. Aligned with its role inhibiting VE cadherin endocytosis, p120-catenin has also been shown to interact with p190RhoGAP at adherens junctions, regulating actin cytoskeleton remodelling via the Rac-dependent antagonism of RhoA [59]. As depleting Nrp1, p120-catenin [10], or Nrp2 in ECs results in junction disassembly and the accumulation of cortical stress fibres, Nrps likely provide the necessary spatial cues to coordinate stress fibre anchorage to junctional sites undergoing nascent assembly and or remodelling. In support of this, it was recently shown that Sema3G, via the Nrp2/plexinD1 signalling pathway, stabilised interactions between β-catenin and VE cadherin to ensure actin cytoskeleton linkage and vascular stability [60]. This hypothesis also runs consistent with our previous published works demonstrating that Nrp2 promotes the activation of the small GTPase Rac1 [11], and the complex association between p120RasGAP and p190RhoGAP to promote polarised integrin recycling [14].

As junctional instability and increased vascular permeability are known to be major risk factors for endothelial activation and dysfunction during the development of atherogenesis, we theorised that Nrp2 may play a protective role against the progression of cardiovascular ischemia. Recent works by Basu *et al*., comparing the transcriptomic profiles of cardiac ECs following ischemic injury support this hypothesis, revealing a significant downregulation in genes regulating EC contraction and vascular tone 2 days (D2) post myocardial infarction (MI). Importantly, ischemic injury at D2 was also found to significantly downregulate Nrp2 gene expression. The authors discuss this early EC response as being defined by the acquisition of pro-inflammatory phenotypes and metabolic changes [61], a commentary that closely mirrors the effects of targeting Nrp2. Indeed, transcriptomic profiling of Nrp2 depleted ECs revealed a notable increase in pro-inflammatory marker expression, including *Il1A*, *Il11*, *Ccl5*, *Cxcl11* and *Cxcl12*, in addition to *Nppb* and *Tnn2*, both predictive biomarkers for cardiac ischemia [26], [28], [31], [37], [39]. Many differentially regulated gene signatures identified in *si*Nrp2 ECs vs Ctrl ECs were also found to correspond with those identified by single-cell sequencing data generated from donors with heart failure compared to healthy tissue, for example *Ankrd1, Nppb*, *Ctgf*, and *Cyr61* [40]. Furthermore, multiple differentially regulated pathways were found to match those identified from our standard RNA-Seq analysis; MAPK signalling (upregulated), Notch signalling and ion channel signalling (downregulated) [40]. A role for Nrp2 regulating chronic inflammatory responses has also been reported in lymphatic ECs (LECs), Mucka *et al*., reporting an increase in tissue inflammation and vascular permeability following Nrp2 depletion [62]. We were surprised however, given this transcriptional shift, not to see changes to the gene expression of major pro-inflammatory cytokines such as Il6, or members of the TGFβ family, which were shown to be upregulated in Nrp1 knockdown ECs [10]. Similarly, no putative interaction between Nrp2 and TGFβR2 was identified by IP LFQ-DIA-LC-MS/MS analysis (as reported between Nrp1 and TGFβR2 [10]), suggesting a degree of disparity between Nrp1 and Nrp2-mediated pro-inflammatory signalling in ECs.

The metabolic switch regulating glycolytic oxidation during EC activation is known to be regulated by hyperlipidaemia and hypoxia in atherosclerotic plaques [41], [43]. Whilst we observed no activation of the HIF1α signalling cascade under basal, normoxic conditions, Nrp2 silencing was found to downregulate genes involved in lipid metabolism, and upregulate genes associated with glycolysis and oxidative stress. Surprisingly, *si*Nrp2 ECs exhibited elevated glycolytic metabolism under basal conditions despite their significantly-induced transcription of *Cited2*, which is known to negatively modulate the hypoxic response during metabolic haemostasis via its competitive binding of the HIF1α subunit [63]. Whilst beyond the scope of this study, we believe that comparing the gene expression profiles of Ctrl and *si*Nrp2 ECs following CoCl_2_ treatment with those cultured under normoxia would provide greater insight into the role of Nrp2 under hypoxic stress.

Of note, it was also recently reported that knockdown of the vasoactive factor angiopoietin-like 4 (Angptl4) in ECs promoted fatty acid uptake and oxidation, in addition to reducing glycolytic metabolism. This was found to result from a direct reduction in Hk2 activity [45]. As Angptl4 was previously shown to complex with both Nrp1 and Nrp2 to promote vascular hyperpermeability [64], together with a significant upregulation of Angptl4 gene expression in *si*Nrp2 ECs, we postulate that a competitive Angptl4-Nrp2 signalling nexus exists to regulate EC activation and metabolic activity under stress.

In summary, we present evidence that Nrp2 functions protectively against adherens junction breakdown in endothelial cells, modulating the transcription of inflammatory genes and EC activation. Our findings therefore suggest that inhibition of endothelial Nrp2 may contribute to the progression of cardiovascular disease in this context.

## Materials and Methods

### Antibodies and reagents

The following primary antibodies were used for experimental analyses:

Nrp2 (SCB: sc-13117, RRID: AB_628044, 1:50; CST: 3366, RRID: AB_2155250, 1:1000), VE cadherin (BD: 555289, RRID:AB_395707, 1:500), β-actin (CST: 8457S, RRID: AB_10950489, 1:2000), P120 catenin (BD: 610134, RRID:AB_397537, 1:100), Phospho-VE cadherin Y685 (Ab119785, RRID:AB_10971838, 1:1000), EEA1 (Ab2900, RRID: AB_2262056, 1:100), Rab7 (CST: 9367, RRID:AB_1904103, 1:100), Lamp1 (CST: 9091, RRID:AB_2687579, 1:100), Ter-119 (R&D: MAB1125, RRID:AB_2297123, 1:100), Fibrinogen β-chain (NB: NBP1-33582, RRID:AB_2293974, 1:100), Nrp1 (R&D: AF566, RRID: AB_355445, 1:100), HIF1-α (Ab179483, RRID:AB_2732807, 1:1000), Hk2 (CST: 2867, RRID:AB_2232946, 1:1000), Endomucin (Sc-65495, SCB; RRID:AB_2100037, 1:500), Vcam1 (Ab134047, RRID:AB_2721053, 1:500), Icam1 (TF: MA5407, RRID:AB_223596, 1:500).

*CST: Cell Signalling Technologies; R&D: R&D Systems; SCB: Santa Cruz Biotechnology; Ab: Abcam; Inv: Invitrogen; NB: Novus Biologicals, BD: BD Biosciences, TF: Thermo Fisher*

### Animal breeding and generation

All experiments were performed in accordance with UK home office regulations and the European Legal Framework for the Protection of Animals used for Scientific Purposes (European Directive 86/609/EEC). All experiments were approved by the Animal Welfare and Ethical Review Board (AWERB) committee at the University of East Anglia, UK. Nrp2 floxed (Nrp2*^flfl^*) mice [65] generated on a C57/BL6 background were purchased from The Jackson Laboratory (Bar Harbour, Maine, USA). Floxed mice were crossed with tamoxifen-inducible PDGFB.iCreER^T2^ mice, provided by Marcus Fruttiger (UCL, London, UK). Gene deletion was achieved via tamoxifen administration *in vivo (see retinal angiogenesis assays and high-fat diet studies)*.

### Cell culture

Primary mouse microvascular lung endothelial cells (mMLECs) were isolated from age-matched (3 – 6 weeks) wild-type (WT) or floxed C57/BL6 mice. Cellular digests were expelled through a 19 gauge-needle and filtered through a 70 µm sterile strainer (Fisher Scientific). Cell pellets were seeded onto plasticware coated with a solution of 0.1 % gelatin containing 10 µg/ml human plasma fibronectin (FN) (Millipore). and collagen type 1. mLMECs were twice positively selected for using endomucin primary antibody and magnetic activated cell sorting (MACS) as previously described by Reynolds & Hodivala-Dilke [66], prior to immortalisation using polyoma-middle-T-antigen (PyMT) as previously described by Robinson *et al*. [67]. ECs were cultured in a 1:1 mix of Ham’s F-12:Dulbecco’s Modified Eagle Medium (DMEM) (low glucose) medium supplemented with 10% Fetal Bovine Serum (FBS), 100 units/mL penicillin/streptomycin (P/S) and 50 μg/mL heparin (Sigma) at 37 °C in a humidified incubator (+ 5% CO_2_) unless otherwise stated. Passaging did not exceed 20. VEGF-stimulation was achieved using 30 ng/ml VEGF-A_164_ (VEGF-A) (murine equivalent to VEGF-A_165_) post 3 hours incubation in serum-free medium (OptiMEM®; Invitrogen). VEGF-A was made in-house as previously described by Krilleke *et al*. [68].

For experimental analyses, plasticware was coated using 10 µg/ml human plasma FN.

To simulate laminar and low shear stress, confluent ECs adhered to coverslips pre-coated with 10 µg/ml FN were placed on an orbital shaker set at 210 rpm for 6 hours at 37 °C in a humidified incubator (+ 5% CO_2_) prior to fixation. ECs adhered to the well periphery (subject to ∼12 dyne/cm^2^) were defined as under laminar flow, whilst ECs adhered at the well centre (subject to ∼5 dyne/cm^2^) were defined as under low shear stress as previously described [69].

Bone marrow-derived monocytic cells were cultured on plasticware at 37 °C in a humidified incubator (+ 5% CO_2_) in Dulbecco’s Modified Eagle Medium (DMEM) (high glucose) medium supplemented with 10% Fetal Bovine Serum (FBS).

### siRNA transfection

ECs were detached using 0.25 % trypsin-EDTA and subject to oligofection using either non-targeting control siRNA (Ctrl) or murine/human-specific siRNA duplexes suspended in nucleofection buffer (200 mM Hepes, 137 mM NaCl, 5 mM KCl, 6 mM D-glucose, and 7 mM Na_2_HPO_4_ in nuclease-free water). Oligofection was performed according to manufacturer’s instructions using the Amaxa 4D-nucleofector system (Lonza).

Mouse-Nrp2 siGENOME siRNA duplexes: D-040423-*03*, D-040423-*04*.

### Co-immunoprecipitation assays

ECs were placed on ice before being lysed at 4 °C in lysis buffer (25 mM Tris-HCl, pH 7.4, 100 mM NaCl, 2 mM MgCl_2_, 1 mM Na_3_VO_4_, 0.5 mM EGTA, 1% Triton X-100, 5 % glycerol, supplemented with Halt^TM^ protease inhibitors) and cleared by centrifugation at 12,000 x g for 20 minutes at 4 °C. Supernatant proteins were then quantified using the BCA assay and equivalent protein concentrations (800 µg) were immunoprecipitated with Dynabeads^TM^ Protein G (Invitrogen) coupled to a primary antibody at 4 °C. Immunoprecipitated proteins were separated by SDS-PAGE and subjected to Western blot analysis.

### Immunoprecipitation data-independent-acquisition label-free quantitative mass spectrometry

Mass spectrometry was carried out by Fingerprints Proteomics Facility, Dundee University, UK, on EC samples immunoprecipitated against Nrp2 primary antibody as described in Co-immunoprecipitation assays. IP’d samples were prepared as described by Benwell *et al*., [14]. Briefly, IP samples were resolved via 1D SDS-PAGE stained with Quick Coomassie Stain. Gel lanes were then excised and subjected to in-gel processing before overnight trypsin digestion. Digested peptides were run on a Q-ExactivePlus (Thermo) instrument coupled to a Dionex Ultimate 3000 HPLC system. Raw data was acquired in Data Independent Acquisition (DIA) mode. Raw data was analysed, and significance was calculated using Spectronaut v17. Gene-Ontology (GO) enrichment analysis was performed using Enrichr software [70], [71], [72]. Protein-protein interaction network maps were visualised using Cytoscape software v3.10.0.

### Western blotting

ECs were lysed in lysis buffer (Tris-HCL: 65 mM pH 7.4, sucrose: 60 mM, 3 % SDS), homogenised and analysed for protein concentration using the bicinchoninic acid (BCA) assay (Pierce). Equivalent protein concentrations were run on 8 % polyacrylamide gels before being subject to SDS-PAGE. Proteins were transferred to a 0.45 µm Amersham Protran® nitrocellulose membrane (GE Healthcare, Amersham) before being incubated in primary antibody resuspended in 5 % milk-powder at 4 °C. Membranes were washed with 0.1 % Tween-20 in PBS (PBST) and incubated in an appropriate horseradish peroxidase (HRP)-conjugated secondary antibody (Dako) diluted 1:2000 for 2 hours at RT. Bands were visualised by incubation with a 1:1 solution of Pierce ECL Western Blotting Substrate (Thermo). Chemiluminescence was detected on a ChemiDoc^TM^ MP Imaging System (BioRad). Densitometric readings of band intensities were obtained using ImageJ^TM^.

### Immunocytochemistry

ECs were seeded onto pre-coated acid-washed, sterilised glass coverslips before fixation in 4 % paraformaldehyde (PFA). Fixed ECs were incubated with 10 % goat serum in PBS 0.3% triton X-100 prior to primary antibody overnight at 4 °C. Following primary antibody incubation, ECs were incubated with an appropriate Alexa fluor secondary antibody diluted 1:200 before mounting using flouromount-G with DAPI^TM^ (Invitrogen). Images were captured using a Zeiss LSM880 Airyscan Confocal microscope with an Axiocam 503 mono camera.

### Retinal angiogenesis assays

Inducible, endothelial specific deletion of Nrp2 was achieved by subcutaneous (SC) tamoxifen injections (50 µl, 2 mg/ml stock) on postnatal (P) days 2-3, followed by intraperitoneal (IP) injections of the same dose on P4-P5. Mice were sacrificed on P6, and retinas harvested. Dissected retinas were fixed in 4 % PFA for 30 minutes before permeabilised in PBS 0.25 % triton-X100. Retinas were then incubated in Dako serum free protein blocking solution for 1 hour, before incubated in primary antibody. Following primary antibody incubation, retinas were washed in 0.1 % triton-X100 and incubated in the appropriate Alexa fluor secondary antibody before being mounted with Flouromount-G. Images were captured using a Zeiss LSM880 Airyscan Confocal microscope with an Axiocam 503 mono camera. All analyses were performed using ImageJ^TM^ software unless otherwise stated.

### RNA library preparation and NovaSeq sequencing

Total RNA was extracted from confluent Ctrl and *si*NRP2 ECs grown under static conditions from 5 independent experiments using the SV Total RNA Isolation System (Promega Cat# Z3100). Standard total RNA sequencing was performed by Genewiz (Azenta Life Sciences) as follows. RNA samples were quantified using Qubit 4.0 Fluorometer (Life Technologies, Carlsbad, CA, USA) and RNA integrity was checked with RNA Kit on Agilent 5300 Fragment Analyzer (Agilent Technologies, Palo Alto, CA, USA). RNA sequencing libraries were prepared using the NEBNext Ultra II RNA Library Prep Kit for Illumina following manufacturer’s instructions (NEB, Ipswich, MA, USA). Briefly, mRNAs were first enriched with Oligo(dT) beads. Enriched mRNAs were fragmented for 15 minutes at 94 °C. First strand and second strand cDNAs were subsequently synthesized. cDNA fragments were end repaired and adenylated at 3’ ends, and universal adapters were ligated to cDNA fragments, followed by index addition and library enrichment by limited-cycle PCR. Sequencing libraries were validated using NGS Kit on the Agilent 5300 Fragment Analyzer (Agilent Technologies, Palo Alto, CA, USA), and quantified by using Qubit 4.0 Fluorometer (Invitrogen, Carlsbad, CA). The sequencing libraries were multiplexed and loaded on the flow cell on the Illumina NovaSeq X Plus instrument according to manufacturer’s instructions. The samples were sequenced using a 2×150 Pair-End (PE) configuration. Image analysis and base calling were conducted by the NovaSeq X Plus Control Software v1.2.0 on the NovaSeq instrument. Raw sequence data generated from Illumina NovaSeq was converted into fastq files and de-multiplexed. One mismatch was allowed for index sequence identification. Enrichment analysis was carried out using Enrichr software. UMAP cluster plot analysis was performed using Appyter.

### ROS assays

ECs were seeded onto pre-coated acid-washed, sterilised glass coverslips overnight at 37 °C. ECs were then incubated with CellROX^®^ Green reagent at a final concentration of 5 µM for 30 minutes at 37 °C, before being washed 3 times with PBS. ECs were the fixed in 4 % PFA for imaging using a Zeiss LSM880 Airyscan Confocal microscope with an Axiocam 503 mono camera.

### Aorta assays

Inducible, endothelial specific deletion of Nrp2 was achieved by IP tamoxifen injections (75 mg/kg bodyweight, 2 mg/ml stock) thrice weekly on 4-week-old mice. Aortas were harvested 25 days following the first tamoxifen injection and fixed in 4 % PFA for 20 minutes. Aortas were then incubated in 30% sucrose/PBS overnight at 4 °C, followed by incubation in a 1:1 mix of 30% sucrose/PBS to OCT. Aortas were then mounted and sectioned in 100 % OCT prior to staining.

### Monocyte attachment assay

Bone marrow-derived monocytes (BMDMs) were generated from flushed bone marrow from tibias and femurs of 6-week-old C57/BL6 WT mice. Isolated bone marrow was treated with red blood cell lysis buffer (RBCLB) for 5 minutes, washed twice with PBS, and cultured for 72 hours. BMDMs were labelled with 1 µM Calcein-AM (Biolegend, Cat# 425201) for 30 minutes. 5×10^5^ labelled BMDMs were incubated with confluent Ctrl and *si*Nrp2 EC-coated coverslips for 1 hour following 6 hours incubation under orbital shear stress (210 rpm). Coverslips were then washed with PBS and fixed in 4% PFA prior to mounting with flouromount-G with DAPI^TM^. Double-positive DAPI-fluorescein BMDM cells/mm^2^ were quantified in a minimum of 15 images per condition per independent experiment.

### Flow cytometry

BMDMs were incubated with confluent Ctrl and *si*Nrp2 ECs for 1 hour under static conditions before being prepared for flow cytometry as previously described by Mckee et al., [73]. Briefly, cells were detached using 0.25 % trypsin-EDTA and directly loaded into wells of a 96-well PCR microplate with fluorescence-minus-one (FMO) controls (prepared using pooled cell suspensions).To prevent non-specific antibody binding, cells were resuspended in TruStain FCX^TM^ anti-mouse CD16/32 antibody (1:200) in fluorescence-activated cell sorting (FACS) buffer (Biolegend, Cat# 101320) at 4 °C. Cells were then incubated in appropriate primary antibody solution, alongside LIVE/DEAD^TM^ Fixable Red (1:400) (Thermo, Cat# L34971) before being fixed in 4 % PFA at 4 °C. AbC^TM^ Total Antibody Compensation Bead Kit (Thermo, Cat# A10497) was used according to manufacturer’s instructions to prepare single stained controls for compensation. All single stained controls, FMOs and samples were run on an LSR-Fortessa^TM^ (BD BioSciences). 10,000 events were recorded for single stained controls and between 100,000 – 200,000 events were recorded for FMOs and samples. FlowJo (v10.9.0, BD) software was used for population gating based on forward and side scatter (FSC, SSC), and fluorescence emissions.

### High-fat diet studies

20-week-old Nrp2*^flfl^* and Nrp2*^flf^*.PDGFB.iCreER^T2^ (Nrp2*^flfl^*.*EC^KO^*) mice received 3 IP injections of 2 mg/ml tamoxifen before being fed a 45% high-fat chow diet for 12 weeks. Aortas were then isolated and stained with Oil-red-O as previously described, [10] to visualise atherosclerotic lesion formation. Briefly, dissected aortas were perfusion fixed in 4 % PFA, washed twice with PBS, then incubated with 60% Oil-red-O solution for 20 minutes. Aortas were then washed once in PBS before mounting and imaging. Atherosclerotic plaque area was determined as % lesion area relative to total aorta area.

## Statistical analysis

All graphic illustrations and analyses to determine statistical significance were generated using GraphPad Prism 9 software and Student’s t-tests unless otherwise stated. Bar charts show mean values with standard error of the mean (+ SEM). Asterisks indicate the statistical significance of p values: p > 0.05 = NS (not significant), *p < 0.05, **p < 0.01, ***p < 0.001 and ****p < 0.0001.

## Data availability statement

The raw data supporting the conclusions of this article will be made available by the authors, without undue reservation, to any qualified researcher.

## Author Contributions

Conceptualisation: CJB, SDR; Formal analyses: CJB, NI, AN, LBV, CAP, LM; Investigation: CJB, NI, AN, LBV, MM, CAP, LM, TL; Resources: SDR; Review and editing: CJB, NI, AN, LBV, MM, CAP, LM, TL, SDR; Visualisation: CJB, Supervision: SDR; Funding acquisition: SDR.

## Acknowledgments

This work was supported by funding from: BHF (grant number PG/22/11033); SDR gratefully acknowledges the support of the Biotechnology and Biological Sciences Research Council (BBSRC); this research was partially funded by the BBSRC Institute Strategic Programme Food Microbiome and Health BB/X011054/1 and its constituent project BBS/E/F/000PR13632. We would like to thank the Quadram Institute Bioinformatics Core and the Quadram Institute Bioimaging Facility at Quadram Institute Bioscience for their expert support in data analysis and imaging, respectively. Their expertise and state-of-the-art resources greatly contributed to this work.

---

**Suppl. Figure 1:**
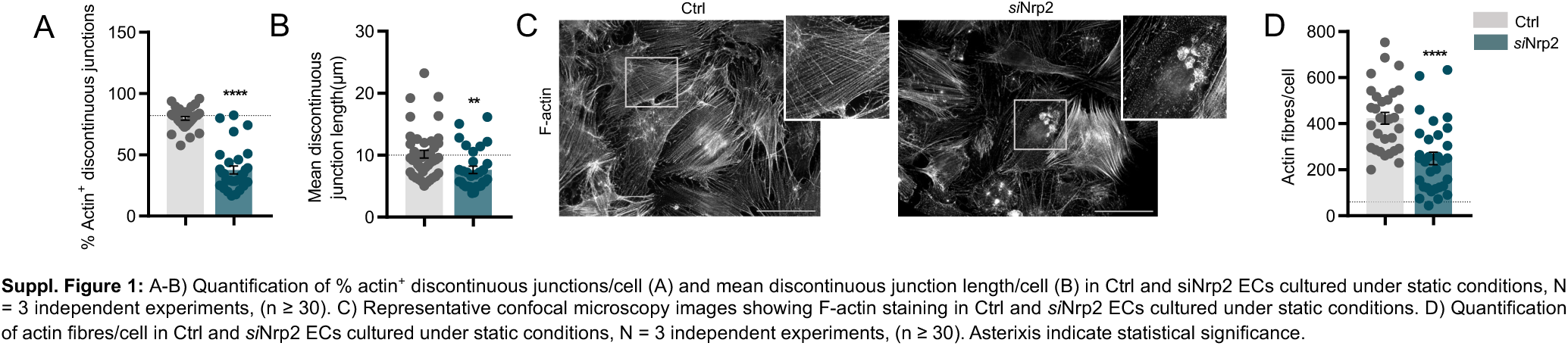
A-B) Quantification of % actin^+^ discontinuous junctions/cell (A) and mean discontinuous junction length/cell (B) in Ctrl and siNrp2 ECs cultured under static conditions, N = 3 independent experiments, (n ≥ 30). C) Representative confocal microscopy images showing F-actin staining in Ctrl and *si*Nrp2 ECs cultured under static conditions. D) Quantification of actin fibres/cell in Ctrl and *si*Nrp2 ECs cultured under static conditions, N = 3 independent experiments, (n ≥ 30). Asterixis indicate statistical significance.

**Suppl. Figure 2:**
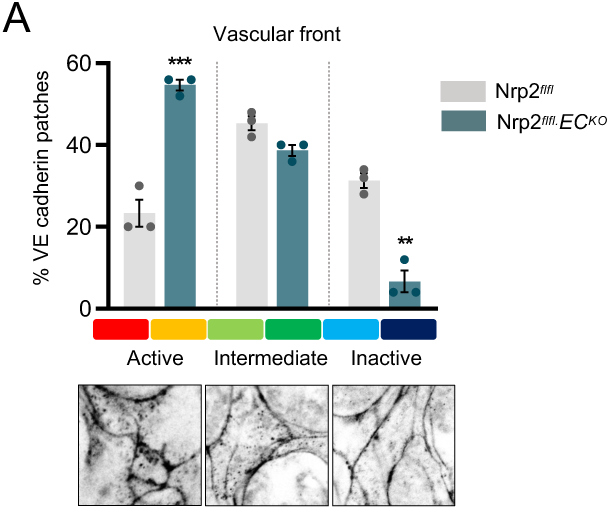
A) Quantification of VE cadherin morphology at the vascular front of Nrp2*^flfl^* and Nrp2*^flfl^*.*EC^KO^* retinas. VE cadherin morphology in each patch/ROI was manually classified using a scale from active/serrated junctions (red), to inactive/continuous junctions (blue). Representative patch images of active, intermediate and inactive VE cadherin morphologies are shown below. N = 3 retinas (> 50 x 100 µm^2^ patches/ROIs quantified per retina). Asterixis indicate significance.

**Suppl. Figure 3:**
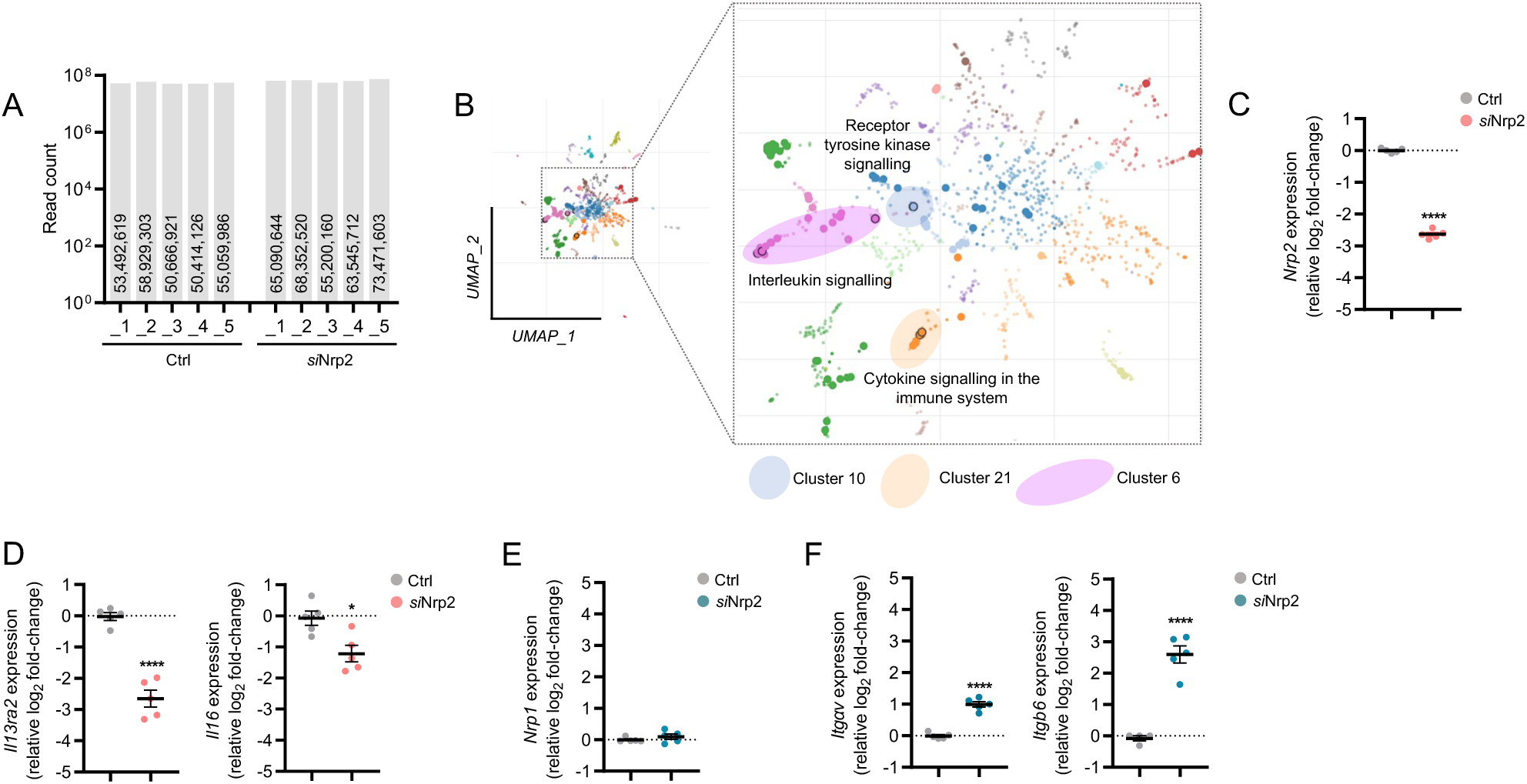
A) Raw gene read counts identified from RNA-seq analysis in Ctrl and siNrp2 samples, N = 5 independent experiments. B) UMAP clustering of upregulated genes following Nrp2 depletion. C) Relative *Nrp2* gene expression in Ctrl and *si*Nrp2 EC RNA-seq samples, N = 5 independent experiments. D) Relative *Il13ra2* and *Il16* gene expression in Ctrl and *si*Nrp2 EC RNA-seq samples, N = 5 independent experiments. E) Relative *Nrp1* gene expression in Ctrl and *si*Nrp2 EC RNA-seq samples, N = 5 independent experiments. F) Relative *αV* and *β6* gene expression in Ctrl and *si*Nrp2 EC RNA-seq samples, N = 5 independent experiments. Asterixis indicate significance.

**Suppl. Figure 4:**
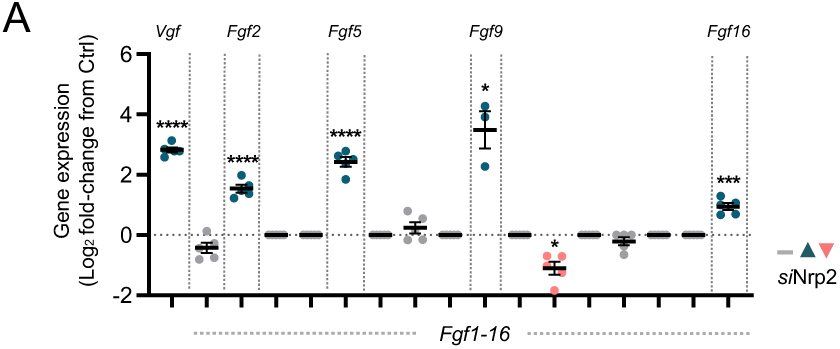
A) Relative gene expression of Fgfs in Ctrl and *si*Nrp2 EC RNA-seq samples, N = 5 independent experiments. Asterixis indicate significance.

**Suppl. Figure 5:**
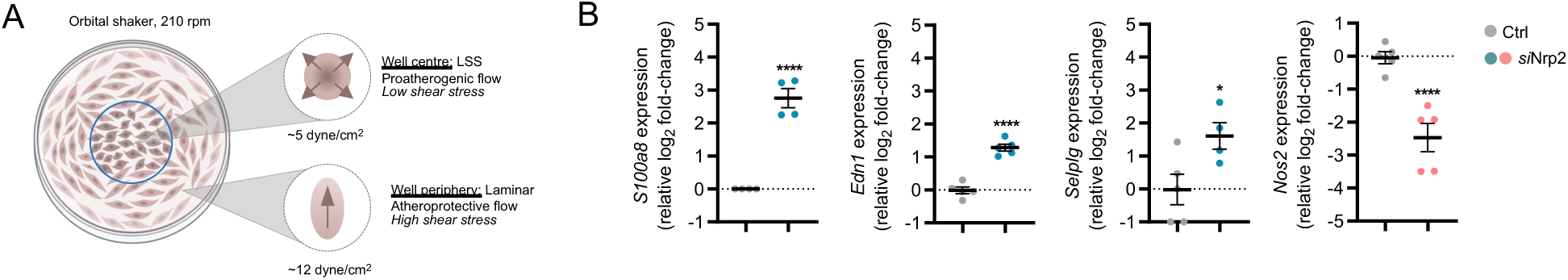
A) Schematic showing orbital shaker method to induce low shear stress (LSS). B) Relative *S100a8, Edn1, Selplg* and *Nos2* gene expression in Ctrl and *si*Nrp2 EC RNA-seq samples, N = 5 independent experiments. Asterixis indicate significance.

## References

[1] E. Dejana, F. Orsenigo, and M. G. Lampugnani, “The role of adherens junctions and VE-cadherin in the control of vascular permeability,” J Cell Sci, vol. 121, no. 13, pp. 2115–2122, Jul. 2008, doi: 10.1242/jcs.017897.

[2] L. Claesson-Welsh, E. Dejana, and D. M. McDonald, “Permeability of the Endothelial Barrier: Identifying and Reconciling Controversies,” Apr. 01, 2021, *Elsevier Ltd*. doi: 10.1016/j.molmed.2020.11.006.

[3] S. Nourshargh and R. Alon, “Leukocyte Migration into Inflamed Tissues,” Nov. 20, 2014, Cell Press. doi: 10.1016/j.immuni.2014.10.008.

[4] W. A. Muller, “How endothelial cells regulate transmigration of leukocytes in the inflammatory response,” American Journal of Pathology, vol. 184, no. 4, pp. 886–896, 2014, doi: 10.1016/j.ajpath.2013.12.033.

[5] B. Engelhardt and H. Wolburg, “Mini review: Transendothelial migration of leukocytes: Through the front door or around the side of the house?,” Nov. 2004. doi: 10.1002/eji.200425327.

[6] D. E. Conway and M. A. Schwartz, “Mechanotransduction of shear stress occurs through changes in ve-cadherin and pecam-1 tension: Implications for cell migration,” Cell Adh Migr, vol. 9, no. 5, pp. 335–339, 2015, doi: 10.4161/19336918.2014.968498.

[7] R. G. Oas, B. A. Nanes, C. C. Esimai, P. A. Vincent, A. J. García, and A. P. Kowalczyk, “P120-catenin and β-catenin differentially regulate cadherin adhesive function,” Mol Biol Cell, vol. 24, no. 6, pp. 704–714, Mar. 2013, doi: 10.1091/mbc.E12-06-0471.

[8] C. M. Chiasson, K. B. Wittich, P. A. Vincent, V. Faundez, and A. P. Kowalczyk, “p120-Catenin Inhibits VE-Cadherin Internalization through a Rho-independent Mechanism,” Mol Biol Cell, vol. 20, pp. 1970–1980, 2009, doi: 10.1091/mbc.E08.

[9] B. A. Nanes et al., “P120-catenin regulates VE-cadherin endocytosis and degradation induced by the Kaposi sarcoma-Associated ubiquitin ligase K5,” Mol Biol Cell, vol. 28, no. 1, pp. 30– 40, Jan. 2017, doi: 10.1091/mbc.E16-06-0459.

[10] E. Bosseboeuf et al., “Neuropilin-1 interacts with VE-cadherin and TGFBR2 to stabilize adherens junctions and prevent activation of endothelium under flow,” Sci Signal, vol. 16, no. 786, 2023, doi: 10.1126/scisignal.abo4863.

[11] A. A. A. Alghamdi, C. J. Benwell, S. J. Atkinson, J. Lambert, R. T. Johnson, and S. D. Robinson, “NRP2 as an Emerging Angiogenic Player; Promoting Endothelial Cell Adhesion and Migration by Regulating Recycling of α5 Integrin,” Front Cell Dev Biol, vol. 8, May 2020, doi: 10.3389/fcell.2020.00395.

[12] B. Favier et al., “Neuropilin-2 interacts with VEGFR-2 and VEGFR-3 and promotes human endothelial cell survival and migration,” Blood, vol. 108, no. 4, pp. 1243–1250, Aug. 2006, doi: 10.1182/blood-2005-11-4447.

[13] N. Gioelli et al., “Neuropilin 1 and its inhibitory ligand mini-tryptophanyl-tRNA synthetase inversely regulate VE-cadherin turnover and vascular permeability,” Nat Commun, vol. 13, no. 1, Dec. 2022, doi: 10.1038/s41467-022-31904-1.

[14] C. J. Benwell, R. T. Johnson, J. A. G. E. Taylor, J. Lambert, and S. D. Robinson, “A proteomics approach to isolating neuropilin-dependent α5 integrin trafficking pathways: neuropilin 1 and 2 co-traffic α5 integrin through endosomal p120RasGAP to promote polarised fibronectin fibrillogenesis in endothelial cells,” Commun Biol, vol. 7, no. 1, Dec. 2024, doi: 10.1038/s42003-024-06320-4.

[15] D. Vestweber, “VE-cadherin: The major endothelial adhesion molecule controlling cellular junctions and blood vessel formation,” Feb. 2008. doi: 10.1161/ATVBAHA.107.158014.

[16] C. M. Warboys, M. Ghim, and P. D. Weinberg, “Understanding mechanobiology in cultured endothelium: A review of the orbital shaker method,” Jun. 01, 2019, Elsevier Ireland Ltd. doi: 10.1016/j.atherosclerosis.2019.04.210.

[17] F. Wessel et al., “Leukocyte extravasation and vascular permeability are each controlled in vivo by different tyrosine residues of VE-cadherin,” Nat Immunol, vol. 15, no. 3, pp. 223–230, Mar. 2014, doi: 10.1038/ni.2824.

[18] C. J. Benwell, J. A. G. E. Taylor, and S. D. Robinson, “Endothelial neuropilin-2 influences angiogenesis by regulating actin pattern development and α5-integrin-p-FAK complex recruitment to assembling adhesion sites,” FASEB Journal, vol. 35, no. 8, Aug. 2021, doi: 10.1096/fj.202100286R.

[19] C. J. Benwell, R. T. Johnson, J. A. G. E. Taylor, C. A. Price, and S. D. Robinson, “Endothelial VEGFR Coreceptors Neuropilin-1 and Neuropilin-2 Are Essential for Tumor Angiogenesis,” Cancer Research Communications, vol. 2, no. 12, pp. 1626–1640, Dec. 2022, doi: 10.1158/2767-9764.crc-22-0250.

[20] A. Fantin et al., “VEGF165-induced vascular permeability requires NRP1 for ABL-mediated SRC family kinase activation,” Journal of Experimental Medicine, vol. 214, no. 4, pp. 1049–1064, Apr. 2017, doi: 10.1084/jem.20160311.

[21] J. Gómez-Escudero et al., “PKM2 regulates endothelial cell junction dynamics and angiogenesis via ATP production,” Sci Rep, vol. 9, no. 1, Dec. 2019, doi: 10.1038/s41598-019-50866-x.

[22] J. Cao et al., “Polarized actin and VE-cadherin dynamics regulate junctional remodelling and cell migration during sprouting angiogenesis,” Nat Commun, vol. 8, no. 1, Dec. 2017, doi: 10.1038/s41467-017-02373-8.

[23] H. Yamamoto et al., “Integrin β1 controls VE-cadherin localization and blood vessel stability,” Nat Commun, vol. 6, Mar. 2015, doi: 10.1038/ncomms7429.

[24] P. Zhu, W. Sun, C. Zhang, Z. Song, and S. Lin, “The role of neuropeptide Y in the pathophysiology of atherosclerotic cardiovascular disease,” Oct. 01, 2016, Elsevier Ireland Ltd. doi: 10.1016/j.ijcard.2016.06.138.

[25] E. Dri et al., “Inflammatory Mediators of Endothelial Dysfunction,” Jun. 01, 2023, MDPI. doi: 10.3390/life13061420.

[26] A. V Sterpetti, “Inflammatory Cytokines and Atherosclerotic Plaque Progression.,” Curr Atheroscler Rep, 2020, doi: 10.1007/s11883-020-00891-3/Published.

[27] L. Cardilo-Reis et al., “Interleukin-13 protects from atherosclerosis and modulates plaque composition by skewing the macrophage phenotype,” EMBO Mol Med, vol. 4, no. 10, pp. 1072–1086, Oct. 2012, doi: 10.1002/emmm.201201374.

[28] A. Zernecke and C. Weber, “Chemokines in atherosclerosis: Proceedings resumed,” 2014, Lippincott Williams and Wilkins. doi: 10.1161/ATVBAHA.113.301655.

[29] M. Lyu et al., “Tnfrsf12a-mediated atherosclerosis signaling and inflammatory response as a common protection mechanism of Shuxuening injection against both myocardial and cerebral ischemia-reperfusion injuries,” Front Pharmacol, vol. 9, no. APR, Apr. 2018, doi: 10.3389/fphar.2018.00312.

[30] G. Nocentini and C. Riccardi, “GITR: A multifaceted regulator of immunity belonging to the tumor necrosis factor receptor superfamily,” Apr. 2005. doi: 10.1002/eji.200425818.

[31] J. Wu, W. Ma, Z. Qiu, and Z. Zhou, “Roles and mechanism of IL-11 in vascular diseases,” 2023, Frontiers Media S.A. doi: 10.3389/fcvm.2023.1171697.

[32] K. C. Wang et al., “Flow-dependent YAP/TAZ activities regulate endothelial phenotypes and atherosclerosis,” Proc Natl Acad Sci U S A, vol. 113, no. 41, pp. 11525–11530, Oct. 2016, doi: 10.1073/pnas.1613121113.

[33] Z. Deng et al., “TGF-β signaling in health, disease, and therapeutics,” Dec. 01, 2024, *Springer Nature*. doi: 10.1038/s41392-024-01764-w.

[34] P. Aluwihare et al., “Mice that lack activity of αvβ6– and αvβ8-integrins reproduce the abnormalities of Tgfb1– and Tgfb3-null mice,” J Cell Sci, vol. 122, no. 2, pp. 227–232, Jan. 2009, doi: 10.1242/jcs.035246.

[35] H. Bräuninger et al., “Matrix metalloproteinases in coronary artery disease and myocardial infarction,” Dec. 01, 2023, Springer Science and Business Media Deutschland GmbH. doi: 10.1007/s00395-023-00987-2.

[36] H. B. Khoukaz et al., “Drug Targeting of Plasminogen Activator Inhibitor-1 Inhibits Metabolic Dysfunction and Atherosclerosis in a Murine Model of Metabolic Syndrome,” Arterioscler Thromb Vasc Biol, vol. 40, no. 6, pp. 1479–1490, Jun. 2020, doi: 10.1161/ATVBAHA.119.313775.

[37] M. Volpe, S. Rubattu, and J. Burnett, “Natriuretic peptides in cardiovascular diseases: Current use and perspectives,” Feb. 2014. doi: 10.1093/eurheartj/eht466.

[38] P. Ponikowski et al., “2016 ESC Guidelines for the diagnosis and treatment of acute and chronic heart failure: The Task Force for the diagnosis and treatment of acute and chronic heart failure of the European Society of Cardiology (ESC). Developed with the special contribution of the Heart Failure Association (HFA) of the ESC,” Eur J Heart Fail, vol. 18, no. 8, pp. 891–975, Aug. 2016, doi: 10.1002/ejhf.592.

[39] A. Shrivastava, T. Haase, T. Zeller, and C. Schulte, “Biomarkers for Heart Failure Prognosis: Proteins, Genetic Scores and Non-coding RNAs,” Nov. 23, 2020, Frontiers Media S.A. doi: 10.3389/fcvm.2020.601364.

[40] A. L. Koenig et al., “Single-cell transcriptomics reveals cell-type-specific diversification in human heart failure,” Nature Cardiovascular Research, vol. 1, no. 3, pp. 263–280, Mar. 2022, doi: 10.1038/s44161-022-00028-6.

[41] I. A. Tamargo, K. I. Baek, Y. Kim, C. Park, and H. Jo, “Flow-induced reprogramming of endothelial cells in atherosclerosis,” Nov. 01, 2023, Nature Research. doi: 10.1038/s41569-023-00883-1.

[42] L. Zhang et al., “STING is a cell-intrinsic metabolic checkpoint restricting aerobic glycolysis by targeting HK2,” Nat Cell Biol, vol. 25, no. 8, pp. 1208–1222, Aug. 2023, doi: 10.1038/s41556-023-01185-x.

[43] L. Li, M. Wang, Q. Ma, J. Ye, and G. Sun, “Role of glycolysis in the development of atherosclerosis,” Aug. 01, 2022, American Physiological Society. doi: 10.1152/ajpcell.00218.2022.

[44] M. Batty, M. R. Bennett, and E. Yu, “The Role of Oxidative Stress in Atherosclerosis,” Dec. 01, 2022, MDPI. doi: 10.3390/cells11233843.

[45] B. Chaube et al., “Suppression of angiopoietin-like 4 reprograms endothelial cell metabolism and inhibits angiogenesis,” Nat Commun, vol. 14, no. 1, Dec. 2023, doi: 10.1038/s41467-023-43900-0.

[46] S. Coma, A. Shimizu, and M. Klagsbrun, “Hypoxia induces tumor and endothelial cell migration in a semaphorin 3F– and VEGF-dependent manner via transcriptional repression of their common receptor neuropilin 2,” Cell Adh Migr, vol. 5, no. 3, pp. 266–275, 2011, doi: 10.4161/cam.5.3.16294.

[47] D. Moris et al., “The role of reactive oxygen species in the pathophysiology of cardiovascular diseases and the clinical significance of myocardial redox,” 2017, AME Publishing Company. doi: 10.21037/atm.2017.06.27.

[48] J.-J. Chiu and S. Chien, “Effects of Disturbed Flow on Vascular Endothelium: Pathophysiological Basis and Clinical Perspectives,” 2011, doi: 10.1152/physrev.00047.2009.-Vascular.

[49] G. Kaplanski et al., “Thrombin-Activated Human Endothelial Cells Support Monocyte Adhesion In Vitro Following Expression of Intercellular Adhesion Molecule-1 (ICAM-1; CD54) and Vascular Cell Adhesion Molecule-1 (VCAM-1; CD106),” 1998.

[50] H. Zhou, C. Zhao, R. Shao, Y. Xu, and W. Zhao, “The functions and regulatory pathways of S100A8/A9 and its receptors in cancers,” 2023, Frontiers Media SA. doi: 10.3389/fphar.2023.1187741.

[51] C. Zouki, C. Baron, A. Fournier, and J. G. nos Filep, “Endothelin-1 enhances neutrophil adhesion to human coronary artery endothelial cells: role of ET A receptors and platelet-activating factor 1,” 1999. [Online]. Available: http://www.stockton-press.co.uk/bjp

[52] P. D. C. Martins et al., “P-selectin glycoprotein ligand-1 is expressed on endothelial cells and mediates monocyte adhesion to activated endothelium,” Arterioscler Thromb Vasc Biol, vol. 27, no. 5, pp. 1023–1029, May 2007, doi: 10.1161/ATVBAHA.107.140442.

[53] S. Hä et al., “Multi-organ expression profiling uncovers a gene module in coronary artery disease involving transendothelial migration of leukocytes and LIM domain binding 2: The Stockholm Atherosclerosis Gene Expression (STAGE) study,” PLoS Genet, vol. 5, no. 12, Dec. 2009, doi: 10.1371/journal.pgen.1000754.

[54] E. D. Muse et al., “A Whole Blood Molecular Signature for Acute Myocardial Infarction,” Sci Rep, vol. 7, no. 1, Dec. 2017, doi: 10.1038/s41598-017-12166-0.

[55] A. Fantin et al., “NRP1 Regulates CDC42 Activation to Promote Filopodia Formation in Endothelial Tip Cells,” Cell Rep, vol. 11, no. 10, pp. 1577–1590, Jun. 2015, doi: 10.1016/j.celrep.2015.05.018.

[56] A. Fantin et al., “NRP1 acts cell autonomously in endothelium to promote tip cell function during sprouting angiogenesis,” 2013, doi: 10.1182/blood-2012-05.

[57] K. Bouvrée et al., “Semaphorin3A, Neuropilin-1, and PlexinA1 Are Required for Lymphatic Valve Formation,” Circ Res, vol. 111, no. 4, 2012, doi: 10.1161/CIRCRESAHA.112.269316/-/DC1.

[58] D. Valdembri et al., “Neuropilin-1/GIPC1 signaling regulates α5β1 integrin traffic and function in endothelial cells,” PLoS Biol, vol. 7, no. 1, 2009, doi: 10.1371/journal.pbio.1000025.

[59] G. A. Wildenberg et al., “p120-Catenin and p190RhoGAP Regulate Cell-Cell Adhesion by Coordinating Antagonism between Rac and Rho,” Cell, vol. 127, no. 5, pp. 1027–1039, Dec. 2006, doi: 10.1016/j.cell.2006.09.046.

[60] D. Y. Chen et al., “Endothelium-derived semaphorin 3G attenuates ischemic retinopathy by coordinating β-catenin-dependent vascular remodeling,” Journal of Clinical Investigation, vol. 131, no. 4, Feb. 2021, doi: 10.1172/JCI135296.

[61] C. Basu, P. L. Cannon, C. P. Awgulewitsch, C. L. Galindo, E. R. Gamazon, and A. K. Hatzopoulos, “Transcriptome analysis of cardiac endothelial cells after myocardial infarction reveals temporal changes and long-term deficits,” Sci Rep, vol. 14, no. 1, Dec. 2024, doi: 10.1038/s41598-024-59155-8.

[62] P. Mucka et al., “Inflammation and Lymphedema Are Exacerbated and Prolonged by Neuropilin 2 Deficiency,” in *American Journal of Pathology*, Elsevier Inc., Nov. 2016, pp. 2803–2812. doi: 10.1016/j.ajpath.2016.07.022.

[63] H. Yoon, J. H. Lim, C. H. Cho, L. E. Huang, and J. W. Park, “CITED2 controls the hypoxic signaling by snatching p300 from the two distinct activation domains of HIF-1α,” Biochim Biophys Acta Mol Cell Res, vol. 1813, no. 12, pp. 2008–2016, Dec. 2011, doi: 10.1016/j.bbamcr.2011.08.018.

[64] A. Sodhi et al., “Angiopoietin-like 4 binds neuropilins and cooperates with VEGF to induce diabetic macular edema,” Journal of Clinical Investigation, vol. 129, no. 11, pp. 4593–4608, Nov. 2019, doi: 10.1172/JCI120879.

[65] A. Walz, I. Rodriguez, and P. Mombaerts, “Aberrant Sensory Innervation of the Olfactory Bulb in Neuropilin-2 Mutant Mice,” 2002.

[66] L. E. Reynolds and K. M. Hodivala-Dilke, “Primary Mouse Endothelial Cell Culture for Assays of Angiogenesis,” in *Breast Cancer Research Protocols*, New Jersey: Humana Press, 2006, pp. 503–510. doi: 10.1385/1-59259-969-9:503.

[67] S. D. Robinson et al., “αvβ3 integrin limits the contribution of neuropilin-1 to vascular endothelial growth factor-induced angiogenesis,” Journal of Biological Chemistry, vol. 284, no. 49, pp. 33966–33981, Dec. 2009, doi: 10.1074/jbc.M109.030700.

[68] D. Krilleke et al., “Molecular mapping and functional characterization of the VEGF164 heparin-binding domain,” Journal of Biological Chemistry, vol. 282, no. 38, pp. 28045–28056, Sep. 2007, doi: 10.1074/jbc.M700319200.

[69] A. Dardik et al., “Differential effects of orbital and laminar shear stress on endothelial cells,” J Vasc Surg, vol. 41, no. 5, pp. 869–880, May 2005, doi: 10.1016/j.jvs.2005.01.020.

[70] Z. Xie et al., “Gene Set Knowledge Discovery with Enrichr,” Curr Protoc, vol. 1, no. 3, Mar. 2021, doi: 10.1002/cpz1.90.

[71] E. Y. Chen et al., “Enrichr: interactive and collaborative HTML5 gene list enrichment analysis tool,” 2013. [Online]. Available: http://amp.pharm.mssm.edu/Enrichr.

[72] M. V. Kuleshov et al., “Enrichr: a comprehensive gene set enrichment analysis web server 2016 update,” Nucleic Acids Res, vol. 44, no. 1, pp. W90–W97, Jul. 2016, doi: 10.1093/nar/gkw377.

[73] A. M. McKee et al., “Antibiotic-induced disturbances of the gut microbiota result in accelerated breast tumor growth,” iScience, vol. 24, no. 9, Sep. 2021, doi: 10.1016/j.isci.2021.103012.

